# Developmental synaptic pruning sharpens neural representations for predictive motor control

**DOI:** 10.1101/2025.01.01.630979

**Authors:** Shivangi Verma, Sarthak Chandra, Shraddha Pathak, Sriram Narayanan, Vatsala Thirumalai

## Abstract

Developmental synaptic pruning is thought to refine neural circuits by eliminating excess connectivity, but whether this process changes the computations performed by developing circuits remains poorly understood. Here, we show that synaptic pruning sharpens neural representations in the developing zebrafish cerebellum. Climbing fiber (CF) inputs to Purkinje neurons (PNs) progressively transition from poly- to predominantly mono-innervation during the first two weeks of life. In vivo whole-cell recordings reveal that individual CFs convey distinct sensorimotor information, and that convergence of differently tuned CFs degrades the precision of PN representations. As this convergence is reduced during development, PN responses become more selective and temporally precise. Later-stage PNs also exhibit stronger predictive responses to expected sensory events and enhanced responses to violations of those expectations, accompanied by improved behavioral adaptation. These findings suggest that developmental synaptic pruning is not simply a process of reducing excess connectivity, but can contribute to the functional specialization of neural circuits by sharpening neuronal representations and enabling increasingly precise predictive sensorimotor computations.

## Introduction

The refinement and maturation of the nervous system rely on effective synaptic pruning (*1–4*). During development, synapses are first overgrown with the help of embryonic cues and then selectively pruned (*5*), with an estimated 50% of synapses being lost during human brain development (*6*, *7*). This phenomenon of synaptic overgrowth and pruning is seen in the peripheral as well as the central nervous systems and across different species (*8*). While the anatomical, molecular and cellular processes driving developmental synaptic pruning have been studied extensively (*9*, *10*), the functional implications of this process for neural computation and behavior remain largely unexplored. Functionally, circuit computational outcomes will depend critically on whether the multiple inputs to a neuron carry similar or different information. If the inputs to a poly-innervated neuron are differently tuned, elimination of all inputs but one will substantially alter neural representations and profoundly influence network outputs and potentially, behavioral outcomes. We test this hypothesis here using the larval zebrafish olivo-cerebellar system.

In the cerebellum, Purkinje neurons (PNs) receive excitatory inputs from parallel fibers (PFs) of granule cells and climbing fiber (CF) axons that originate from neurons of the inferior olivary nucleus. The CF to PN synapse is one of the strongest excitatory synapses known in vertebrates and triggers a large calcium influx into PNs (*11*, *12*). At birth, every PN in mammals receives input from multiple CFs (poly-innervation), that are then pruned during post-natal life such that each PN is innervated by only a single CF (mono-innervation). The transition of mammalian PNs from poly- to mono-innervation is a closely orchestrated activity-dependent process involving the CF-PN and PF-PN synapses, metabotropic glutamate receptors, voltage-dependent calcium channels and several retrograde and anterograde signaling pathways (*13*, *14*). These processes help in maintaining and strengthening a single CF input while weakening and ultimately removing all other CF inputs.

Classical cerebellar learning models have underscored the role of CFs as the teaching signal in motor learning (*15–17*). More recently, studies have shown that CFs encode motor kinematics, sensory variables, and errors in sensory predictions, decisions, and rewards during a variety of learning tasks (*18–25*). Recently, we showed that CFs encode predictions and prediction errors which enable faster behavioral responses (*22*). We asked whether individual CFs carry distinct sensorimotor information, and if so, how does developmental selection among competing CF inputs alter the information represented by their target PNs.

In mammals, CF synapse elimination occurs during stages when motor behaviors are largely absent. This precludes our ability to understand the effect of CF synapse elimination on cerebellar computation and cerebellum dependent behaviors. In contrast, zebrafish larvae become independent organisms, navigating through complex aqueous environments while their cerebellum is still undergoing maturation (*26*, *27*). Further, zebrafish, with its conserved cerebellar circuitry (*28*), allow us to observe developmental circuit refinement and its consequences for sensorimotor computation and behavior in the same animal.

Here, we asked whether developmental elimination of CF inputs changes the sensorimotor representations in PNs and their capacity to process predictable and unexpected sensory events. Using the larval zebrafish, where cerebellar maturation occurs alongside active sensorimotor behavior, we combined functional measurements of CF innervation with in vivo electrophysiology, population calcium imaging, behavioral assays, and computational modeling. We find that developmental refinement from poly- to predominantly mono-innervation is accompanied by increased selectivity and temporal precision of Purkinje neuron representations. Individual CFs convey distinct sensorimotor information, such that their convergence reduces the precision of the resulting Purkinje neuron representation. As this convergence is reduced during development, Purkinje neurons increasingly express predictive responses to expected sensory events and responses to violations of those expectations, accompanied by improved behavioral adaptation. Together, these findings suggest that developmental synaptic pruning can contribute to the functional specialization of neural circuits by selectively eliminating competing inputs and sharpening the representations that support predictive sensorimotor computations.

## Results

### CF inputs to PNs undergo developmental changes consistent with pruning

Cerebellar PNs receive contralateral inputs from the inferior olivary nucleus via climbing fibers (CFs) (Fig. 1A). We evaluated CF inputs to PNs during the first two weeks of development, focusing on early (4-5 days post fertilization or dpf) and late (11-14 dpf; up to 19 dpf for fictive swim experiments) larval stages, while assessing intermediate stages (7-8 dpf) in some experiments (Fig. 1B). CF inputs to PNs in larval zebrafish can be easily distinguished in whole-cell voltage-clamp and current-clamp recordings as well as loose-patch extracellular recordings due to their large amplitude and stereotypical kinetics (*29*) (Fig. 1C). We recorded spontaneous excitatory post-synaptic currents (EPSCs) from PNs in awake larval zebrafish in early and late stages and found that CF inputs in late-stage larvae have reduced frequency compared to early stages, as reported in our previous study as well (*29*) (Fig. 1C, D). Further, we also observed that the CF EPSCs in the late stages were less variable in amplitude and exhibited lower coefficient of variation (CV) compared to the early stages (Fig. 1C, E). We hypothesized that these results could be explained by (i) synaptic pruning, (ii) changes in short term synaptic plasticity of the CF-PN synapses or (iii) developmental alterations in CF firing rates.

**Figure 1.**
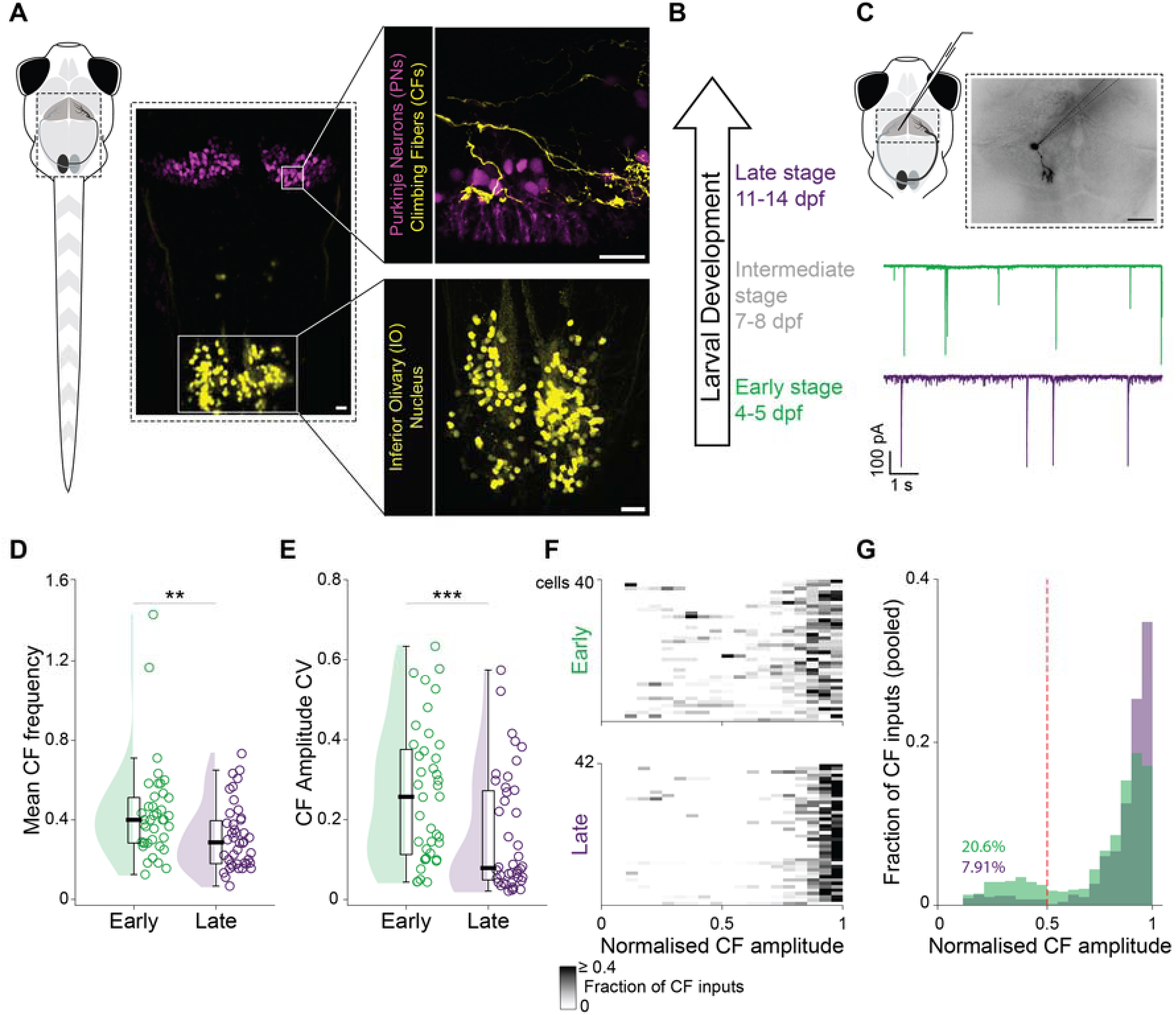
CF-PN synapses show signatures of pruning during first two weeks of larval development in zebrafish. **A.** Schematic of the olivo-cerebellar circuit (highlighted) in larval zebrafish. Inset: Inferior olivary neurons and their climbing fiber (CF) projections (yellow, labelled in a transgenic hspGFFDMC28C:gal4, UAS:gfp larva) superimposed with cerebellar Purkinje neurons (magenta), labelled using Ca8-cfos:ecfp/dendra2). Scale bars 20 µm. **B.** Developmental stages investigated in the current study: 4-5 dpf (also referred to as early larval stage, green), 7-8 dpf (also referred to as intermediate larval stage, grey) and 11-14 dpf (also referred to as late larval stage, purple). **C.** Top: Schematic of *in vivo* whole-cell patch-clamp recording in PNs. Inset: PN filled with sulforhodamine after the recording. Scale bar 20 µm. Bottom: Representative voltage-clamp recordings of spontaneous CF excitatory postsynaptic currents (EPSCs) from a single PN from an early (top, green) and late (bottom, purple) stage larva. **D.** Reduction in spontaneous CF input frequency (Hz) from early to late developmental stages. Median ± IQR : 0.4 ± 0.229 (early), 0.286 ± 0.216 (late) (Mann-Whitney U test, P = 0.006, Cohen’s d = 0.586 ). **E.** Reduction in coefficient of variation (CV) of CF EPSC amplitudes. Median ± IQR : 0.256 ± 0.262 (early), 0.078 ± 0.223 (late) (Mann-Whitney U test, P = 0.0009, Cohen’s d = 0.644). **F.** Heatmap showing the spread of CF EPSC amplitudes (normalised to largest amplitude recorded for each PN). Each row shows the fractional distribution of normalized EPSC amplitudes from one cell. **G.** Reduction in CF inputs with amplitudes less than half of the maximum amplitude from early (20.631%) to late (7.91%) developmental stage (pooled data from panel F). Red dashed line indicates half of the maximum amplitude. Holding potential: -55mV. Recording duration 45 - 180 s. N = 40 and 42 cells (from 34 and 40 larvae) for early and late stages, respectively. Colours *green* and *purple* represent early (4-5 dpf) and late (11-14 dpf) larval stages, respectively. **P < 0.01, ***P<0.001.

If synaptic pruning underlies these changes, the higher frequency of CF inputs seen in PNs at the early stage could be due to poly-innervation by multiple CFs and the varying synaptic strength between synapses formed by multiple fibers can explain the larger variation in EPSC amplitudes observed in early stage PNs. CF-PN synaptic pruning in mammals proceeds via the selective strengthening and stabilization of a single fiber and progressive weakening and elimination of all the others (*9*). Consistent with this view, when we normalized EPSC amplitudes to the maximal amplitude within each cell, we found that in early-stage larvae, EPSCs were of small, medium and large amplitudes, whereas at later stages, large amplitudes predominated (Fig. 1F). The distribution of amplitudes changed such that in early-stage larvae, 20.63% of the pooled amplitudes were less than half of the peak value while in late-stage larvae, this fraction dropped to 7.91% (Fig. 1G). In addition, the EPSC amplitudes were found to be significantly higher in the late larval stage, although with small effect sizes (Fig. S1A). These results point us towards developmental pruning of CF to PN synapses. Nevertheless, we considered the other two alternate possibilities as well.

The CF to PN synapse shows short term depression due to its high release probability (*30–32*). It is possible that the developmental changes in EPSC amplitude variability arise from changes in short term plasticity at these synapses. We probed this angle by only considering CF EPSCs that were spaced in time by more than 200 ms for our analysis. The increase in CF EPSC amplitudes and the decrease in CV of amplitudes persisted even after applying this rule (Fig. S1B, C). Next, we tested this possibility directly by measuring the paired pulse depression (PPD) at these synapses across these developmental stages. We stimulated CFs with two maximal intensity current pulses at interstimulus intervals (ISIs) of 30, 35 and 50 ms, while recording evoked EPSCs from PNs (Fig. S2A). At these ISIs, the paired pulse ratio (see Methods) is less than 1, indicating short term depression at the CF to PN synapse (*32*). While the PPR increased with increasing ISIs as would be expected, we could not detect any differences that might be explained due to developmental stage (Fig. S2B).

The decrease in CF EPSC frequency might also result from changes in the spontaneous firing rates of inferior olivary neurons. Using loose-patch electrophysiology, we recorded spontaneous spikes from the somata of GFP-positive olivary neurons in semi-intact ventral-up preparations of transgenic hspGFFDMC28C:Gal4; UAS-GFP larvae at early and late stages (Fig. S2C, see Methods). The average spontaneous firing rates of olivary neurons showed no significant differences between early and late larval stages (Fig. S2D).

Taken together, the above results suggest that PNs are poly-innervated by CFs in early-stage larvae and are likely to get pruned to mono-innervation by later stages, consistent with similar suggestions from previous studies (*29*, *33*).

### Functional confirmation of CF to PN synaptic pruning

Upon evaluating cerebellar PNs in EM connectome of a 7dpf larval zebrafish (*34*), we found PNs with multiple CF innervations (Fig. 2A). This anatomical evidence suggests occurrence of evolutionarily conserved poly-innervation of PNs by CFs during early larval development. To ascertain whether multiple CFs form functional synapses with PNs in larval zebrafish and that PNs proceed from poly-innervation by CFs to mono-innervation, it is necessary to directly assess the innervation status at various stages. To do this, we performed field stimulation of CFs with graded current intensities and simultaneously recorded evoked EPSCs from PNs using whole-cell voltage-clamp (Fig. 2B) (*4*). Brief current pulses were delivered to create a local electric field at the entry point of CFs into the cerebellum (Fig. 2B). A full range of current intensities from 100% failure to maximal EPSC amplitude in all trials was tested (Fig. 2C). At low intensities, stochastic stimulation of single CFs allowed recruitment of individual fibers. As a result, a graded response was seen in the form of multiple discrete amplitudes of CF EPSCs (also referred to as CF steps), corresponding to the different synaptic strengths of the innervating fibers (Fig. 2C, top). Whereas, in mono-innervated PNs, the responses were either failures or EPSCs of a single amplitude (single step) (Fig. 2C, bottom). At high intensities, a single step was seen due to maximal stimulation of the CFs in both poly- and mono-innervated PNs (Fig. 2C).

**Figure 2.**
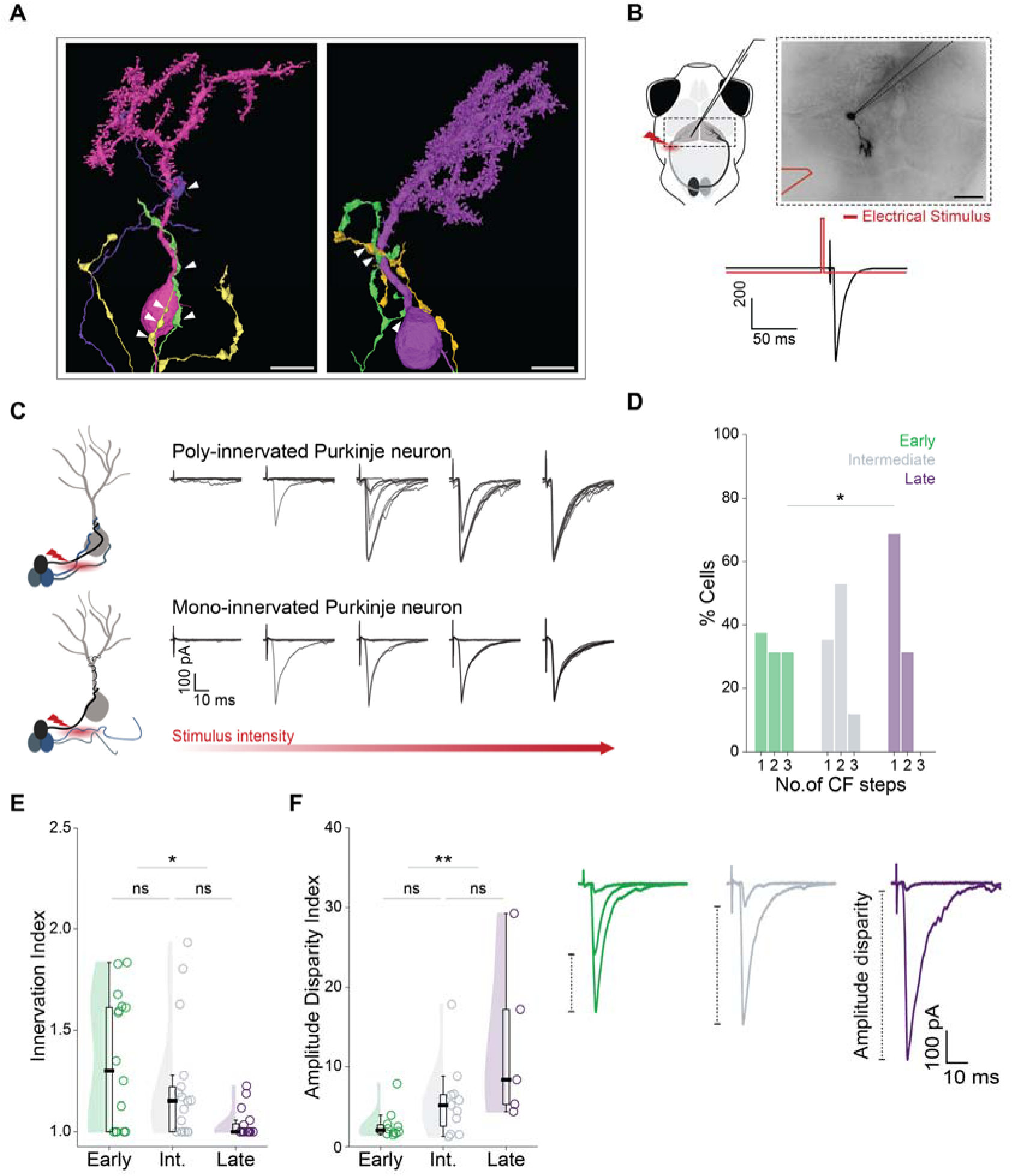
PNs transition from poly-innervation to mono-innervation by CFs during the first two weeks of life. **A.** Electron microscopy reconstructions (*from Petkova et al., 2025* (*34*)) highlighting anatomical evidence of poly-innervated PNs in 7dpf larva. PN with 3 (left) and 2 (right) CF innervations. White arrows indicate synaptic contacts between CFs (yellow, green, purple) and PN (magenta/purple). Scale bars 5 µm. **B.** Top: Schematic of electrophysiological assay for evaluating innervation status. PNs recorded *in vivo* in whole-cell patch-clamp mode with simultaneous electrical stimulation of climbing fibers (**⚡**). Inset: A patched PN with CF stimulation using bipolar electrode (red) placed at the edge of the cerebellum. Bottom: Evoked CF EPSC (black) recorded from a PN after stimulating with a 100 µs long current pulse (red). **C.** Representative traces of evoked CF EPSCs recorded from a poly-innervated PN with multiple discrete amplitude steps (top) and a mono-innervated PN with single amplitude step (bottom). Superimposed trials are shown for each intensity. Graded current intensities between 10 - 480 µA were used. For each intensity, a series of ten pulses, 100 µs long, were delivered at a frequency 0.2 Hz. **D.** Discrete frequency distribution histograms for evoked CF EPSC amplitude steps recorded from PNs over development (One-way ANOVA, P = 0.035). **E.** Reduction in innervation index over early development. Median ± IQR : 1.299 ± 0.613 (early), 1.152 ± 0.221 (intermediate), 1 ± 0.04 (late) (Kruskal Wallis test for multiple comparisons, P = 0.02) followed by post hoc pairwise Mann-Whitney U test, with Holm-Bonferroni correction for early vs intermediate (P = 0.472, Cohen’s d = 0.402), intermediate vs late (P = 0.051, Cohen’s d = 0.829), early vs late (P = 0.031, Cohen’s d = 1.273)). **F.** Left: Developmental increase in amplitude disparity index for evoked CF EPSCs in poly-innervated PNs (shown in panel D). Right: Representative traces showing differential amplitude disparity between evoked CF EPSCs in poly-innervated neurons. Median ± IQR : 2.112 ± 1.062 (early), 5.242 ± 3.914 (intermediate), 8.396 ± 11.878 (late) (Kruskal Wallis test for multiple comparisons (P = 0.022) followed by pairwise Mann-Whitney U test with Holm-Bonferroni correction for early vs intermediate (P = 0.339, Cohen’s d = 0.809), intermediate vs late (P = 0.339, Cohen’s d = 1.049), early vs late (P = 0.007, Cohen’s d = 1.685)). Holding potential: -55mV. N = 16, 17, 16 cells (from 11, 17, 13 larvae) for early, intermediate, late stages, respectively. Colours *green*, *grey* and *purple* represent early (4-5 dpf), intermediate (7-8 dpf) and late (11-14 dpf) larval stages, respectively. *P < 0.05, **P < 0.01, ns P > 0.05.

Upon sampling across larval development, we found a mixed distribution of poly- and mono-innervated PNs, with an increase in mono-innervation over development (Fig. 2D). During the early larval stage, two or more CF steps (poly-innervation) were seen in 10 out of 16 PNs (62.5%), compared to 11 out of 17 PNs (64.5%) in the intermediate larval stage and 5 out of 16 PNs (31.25%) in the late larval stage (Fig. 2D). While the number of steps represents an absolute measure of CFs associated with a PN, it doesn’t account for the varying strengths with which individual fibers are connected to poly-innervated PNs. We used two metrics to quantify the relative strengths of the innervating CFs-innervation index and amplitude disparity index (see Methods section). The ‘innervation index’ is an inverse measure of the proportional contribution of the maximal step CF amplitude to the sum of all step amplitudes. It takes a value equal to 1 for mono-innervated PNs and higher values for poly-innervated PNs. In poly-innervated PNs, with a dominant CF and other weaker inputs, the innervation index takes values close to 1. In cases where the inputs have comparable strengths, the innervation index takes values that are higher than 1. We observed that the innervation indices decline from early (1.299 ± 0.613, Median ± IQR) to intermediate larval stage (1.152 ± 0.221) with an effect size of 0.402 (Fig. 2E). Notably, a shrinkage of 63.94% in the interquartile range during this period brings the intermediate innervation indices closer to 1 (Fig. 2E). This decline continues into the late stage (1 ± 0.04), resulting in a significant reduction from early larval stage with an effect size of 1.273 (Fig. 2E). We used a second measure, the amplitude disparity index to ask how different the largest and the second largest EPSC amplitudes were in PNs that had at least 2 steps (see

Methods). As expected, the amplitude disparity index increased from early to late stages demonstrating that in late-stage larvae, those PNs that had 2 steps, had one dominant CF input, while the other fiber was severely weakened (Fig. 2F).

Based on the above, we can conclude that at early larval stages, PNs are predominantly poly-innervated by CFs of comparable strengths, while at late larval stages, PNs are predominantly mono-innervated. Even when innervated by more than one CF in late larval stage, there is one strong input with the other(s) being relatively weak and less influential.

### Pruning affects the precision, salience and latency of sensori-motor errors

Zebrafish larvae stabilize themselves in slowly moving water bodies by generating swims to compensate for optic flow, in a behavior called the optomotor response (OMR) (*35–37*). In head-restrained larvae, OMR can be reliably induced by presenting black and white gratings and coupling grating movement to larval swims in closed loop (*22*, *38*). When grating movement is uncoupled from larval swims (open loop), larvae sense the mismatch between their visual sensory input and motor output and cease to generate swims for several minutes (*39*). CFs are known to carry error signals when animals encounter a sensori-motor mismatch (*40–43*).

As a first step towards understanding the functional relevance of pruning, we induced a sensori-motor error by presenting ‘open loop’ optic flow that lacks visual feedback (*27*). As the larvae engaged with the optic flow, we recorded the CF inputs from single PNs using whole-cell patch-clamp alongside fictive swims from axial motoneuronal axons (Fig. 3A). When larvae detect that their motor output is ineffective (open loop), they adjust bout parameters even within an ongoing swim bout (*27*), approximately within 300 ms of bout start. Therefore, we looked at this time window relative to bout start to ask whether CF inputs are differentially engaged during different developmental stages.

**Figure 3.**
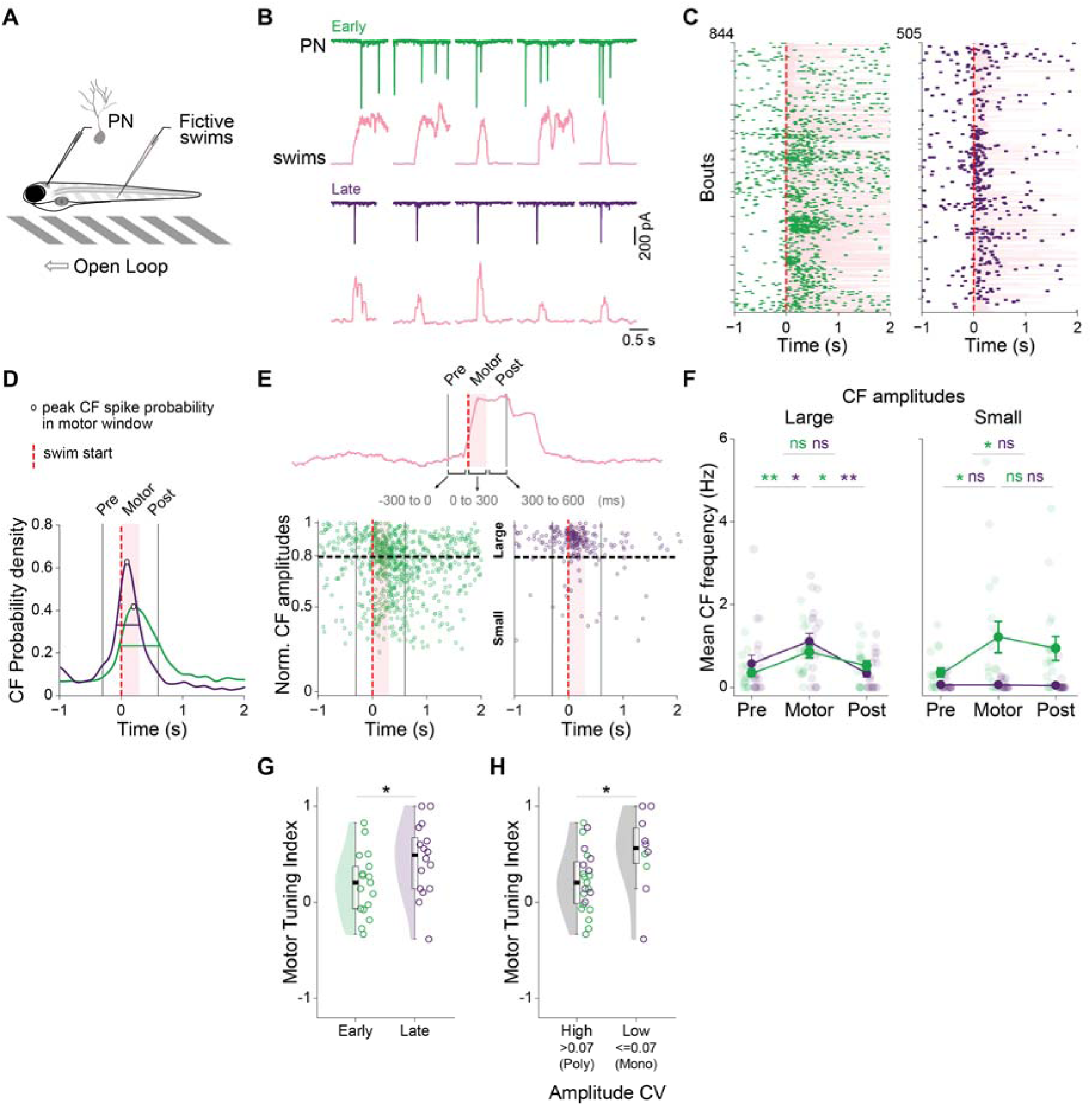
Improved fidelity and temporal precision of sensori-motor mismatch signals in PNs after synaptic pruning. **A.** Schematic of fictive motor recording combined with *in vivo* whole-cell patch-clamp recordings from PNs during optic flow (open loop) presentation. Optic flow consisted of seven trials, each with 10 s stationary followed by 10 s forward-moving gratings. The experiment was performed using current-clamp first and then repeated with voltage-clamp (Holding potential: -55mV) for each cell. Amplitude analysis was performed using voltage-clamp data, while frequency analysis included both current-clamp and voltage-clamp recordings. **B.** Representative traces of voltage-clamp recordings from a PN along with simultaneous fictive swim bouts (pink) in early (top, green) and late (bottom, purple) stage larvae. **C.** Raster plots showing CF inputs around the onset of swim bouts in early (left) and late (right) stage larvae. Dashed red line at 0 marks the start of swim bouts. Ticks on y-axis separate different PNs. Each row is one swim bout. Duration of swim bouts (up to 2s) is shaded in pink. **D.** Gaussian Kernel Density Estimate (KDE) of CF inputs around swim bout onset (red dashed line), with data pooled from all swim bouts shown in panel C (also see Fig. S3B). Open circles and horizontal lines in the respective KDE plots indicate peak CF density and full width at half maximum. Shaded pink region is the motor window and vertical black lines represent the start and end of the pre and post motor window, respectively (described further in panel E (top)). **E.** Top: Time around swim onset is divided into pre (-300 ms to 0 ms), motor (0 ms to 300 ms), and post (300 ms to 600 ms) windows to further evaluate the nature of CF inputs associated with motor error. Bottom: Scatter plot showing time of occurrence of CF EPSCs versus their normalized amplitudes. CF EPSCs were categorized as large or small using a threshold of 0.8 (black dashed line), based on the amplitude distribution in Fig. 1F. Vertical black lines mark the start and end of the pre and post motor window, respectively and the pink region indicates the motor window. **F.** Left: Large CF EPSCs exhibit a motor window specific increase in frequency (Hz) for both developmental groups. Mean ± SEM (pre, motor, post): 0.354 ± 0.09, 0.855 ± 0.137, 0.537 ± 0.11 (early); 0.579 ± 0.196, 1.101 ± 0.2, 0.342 ± 0.095 (late). Right: Sustained elevation in the frequency of small CF EPSCs beyond the motor window during the early larval stage. Mean ± SEM (pre, motor, post): 0.362 ± 0.109, 1.212 ± 0.38, 0.939 ± 0.284 (early); 0.067 ± 0.032, 0.067 ± 0.024, 0.052 ± 0.041 (late) (Friedman test for multiple comparisons, P = 0.0167, 0.002 (large amplitudes for early and late stages, respectively), 0.039, 0.676 (small amplitudes for early and late stages, respectively). Post hoc pairwise comparisons were performed using one-sided Wilcoxon signed-rank test with Holm-Bonferroni correction to test the increase in CF frequency from pre to motor and post windows, and the decrease in CF frequency from the motor to post window). **G.** Motor tuning indices for early and late stage PNs. Median ± IQR : 0.206 ± 0.441 (early), 0.489 ± 0.522 (late) (independent t-test, P = 0.049, Cohen’s d = 0.71). **H.** Motor tuning indices for PNs with high (poly-innervated) and low (mono-innervated) coefficient of variation (CV) of CF EPSC amplitudes. Median CV observed at late stage (∼0.07) was used as threshold for classification. An upward shift can be seen in the Motor tuning index of PNs with low CV. Median ± IQR : 0.206 ± 0.436 (low CV), 0.561 ± 0.366 (high CV) (independent t-test, P = 0.032, Cohen’s d = 0.849). N = 17, 19 cells (from 16, 19 larvae) for early and late stages, respectively. Colours *green* and *purple* represent early (4-5 dpf) and late (15-19 dpf) larval stages, respectively. Red dashed line indicates the onset of swim bouts. *P < 0.05, **P < 0.01, ns P > 0.05.

We observed an increase in the frequency of CF inputs around initiation of swim bouts in both early and late larval stages (Fig. 3B, C). Upon examining individual swim bouts, we found 33.33% ± 26.72 (early stage) and 26.47% ± 34.145 (late stage) of them were accompanied by at least one CF input within 300 ms window from swim start, which we refer to as the motor window (Fig. S3A). The late stage PNs see a peak CF occurrence probability at 95 ms, which is a physiologically relevant latency for cerebellar plasticity and motor learning (*44–46*). However, in early stage PNs, this peak occurs with a much longer delay of 205 ms from swim onset (Fig. 3D). Additionally, late stage PNs also showed a temporally locked elevation in frequency with a peak full width at half max of 380 ms, compared to 656 ms in early stage PNs (Fig. 3D).

Therefore, CF EPSCs exhibit temporal precision and sharper tuning to swim start times in late stage larvae. Furthermore, peak CF probability around motor start increased by 50.388% in the late stage, suggesting an improved salience of motor error (Fig. 3D). Collectively, these results suggest that when there is sensori-motor mismatch leading to an olivary error signal, the error is conveyed with shorter latency, greater temporal precision and higher salience in the later stage mono-innervated PNs compared to the early-stage poly-innervated PNs (Fig. 3D, S3B).

Since there are differences in basal CF frequencies between early and late stages, we compared the specific modulation of CF inputs during the motor window relative to comparable time windows before and after bout start within the respective stages (Fig. 3E, top). We normalized CF EPSCs to their maximal amplitudes and used a threshold of 0.8 to designate events as large or small (Fig. 3E, bottom). In both early- and late-stage larvae, there was a significant up-modulation of large amplitude CF EPSCs in the motor window compared to the time windows before or after it, of similar durations (Fig. 3F, left). However, in the case of the small amplitude events, late-stage larvae showed no modulation while early-stage larvae showed an increase in the frequency of these events in the motor window and in the period following it (3F, right). This suggests that sharper motor tuning of CF inputs in late-stage perhaps arises from the emergence of a large (dominant) CF input and the elimination of weaker inputs.

Early-stage larvae tend to have bouts that are longer in duration compared to late-stage larvae (Fig. S3C, (*47*)). We wondered if in the early-stage larvae, CF EPSCs arrived later with respect to bout start but well within the ongoing motor bout. In this case, one can expect that early-stage larvae might have a reduced number of CF EPSCs during the first 300 ms of the bout (“motor” window) and that there will be differences in the distribution of CF EPSC timing with respect to bout start. We found no differences in the percent bouts with at least one CF EPSC in the motor window in early- and late-stage larvae (Fig. S3A). We also found no evidence that the timing of CF EPSCs within the motor window was dependent on bout duration or that it was developmentally altered (Fig. S3D). These results indicate that the developmental changes in bout duration do not contribute to the alterations in motor error encoding we observed.

Next, we asked whether the observed relationship between CF innervation and motor tuning reflects causation rather than correlation. To determine whether mono-innervated PNs are more strongly tuned to motor activity, we calculated the ‘motor tuning index’ as a measure of specific modulation of CF inputs within the motor window. This was calculated as the difference between CF frequency in the motor window and mean of pre and post motor CF frequencies, over their sum (see Methods). We found that the motor tuning indices in late-stage PNs were larger than those of early-stage PNs (Fig. 3G). We then classified the PNs with high CV of CF EPSC amplitudes as poly-innervated, and those with low CV as mono-innervated (Fig. 3H) based on Fig. 1E. The resultant groups of mono-innervated PNs showed a significantly higher motor tuning index (Fig. 3H).

From the above, we can conclude that in early-stage larvae, due to the occurrence of many CF inputs of diverse strengths, the temporal precision, salience and latency of reporting a sensori-motor mismatch are compromised. As the PNs transition to mono-innervation by late larval stages, these parameters of error encoding improve.

### Late stage PNs receive direction-selective CF inputs

The optomotor stimulus is directional and therefore, any error conveyed to PNs should also be directional if error correction is to be achieved. Indeed, it is known that in mammals complex spikes in Purkinje neurons, triggered by olivary inputs are directionally tuned (*44*, *48*, *49*).

Recently, such direction selectivity in visual input was also shown in zebrafish inferior olivary neurons (*19*). We wondered if the deficits in error encoding of the early-stage larvae during open loop OMR could result from poor directional tuning of CF inputs to PNs. We hypothesized that when larvae encounter optic flow in different directions, a poly-innervated PN would likely receive CF inputs for multiple directions, due to convergence of more than one direction selective CF. In contrast, a mono-innervated PN would receive a single, direction-specific CF input. To test this, we presented optic flow in eight different directions and simultaneously recorded CF inputs from single PNs using loose-patch electrophysiology (Fig. 4A). Over development, we identified mixed PN populations receiving varying degrees of direction-selective CF inputs, ranging from highly selective to completely non-selective (Fig. 4B). We computed a preference index (PI) for each direction (Fig. 4C) (see Methods) and a direction selectivity index (DSI) for every PN as the vector sum of PIs (corresponding to the eight directions) divided by the sum of the magnitude of PIs (see Materials and Methods). A PN receiving CF inputs only for a single direction will have a DSI of 1, whereas if it receives CF inputs equally for all eight directions, the DSI will be 0. We found a mixed distribution of DSIs over development, with a gradual shift towards 1 (Fig. 4D, E). Late-stage PNs showed significantly higher DSIs as compared to early stage with an effect size of 0.96 (Fig. 4E). These results are consistent with the idea that in early-stage larvae, directional error information from the inferior olive is corrupted by several CFs innervating a single PN, leading to less precise representations of error.

**Figure 4.**
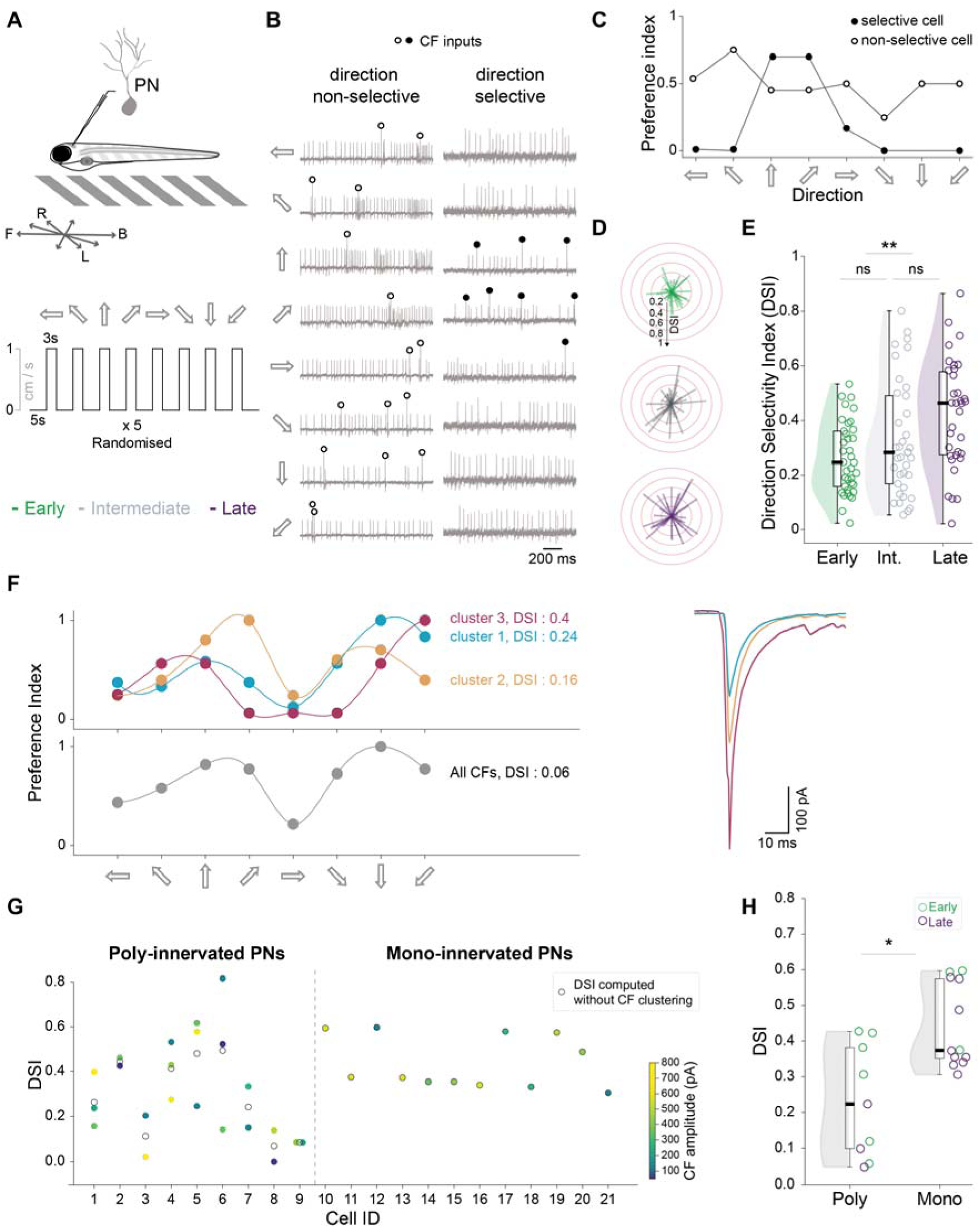
CF inputs to PNs show greater directional selectivity to visual stimuli during late larval stages. **A.** Top: Schematic of loose-patch recording from PNs during optic flow (open loop) presentation in eight different directions-left, forward-left, forward, forward-right, right, backward-right, backward, backward left. Bottom: Trial structure for directional stimulus. Directional optic flow was organized into five blocks, with each block containing eight trials, each corresponding to one of the eight directions chosen randomly. Trials consisted of 5 s stationary followed by 3 s moving gratings. Direction was switched at the end of the moving phase of each trial. **B.** Representative loose-patch recordings from PNs 1.5 s after flow onset. Two PNs are shown-one with direction non-selective CF inputs (left, open circles) and the other with direction selective CF inputs (right, filled circles). These recordings were from 5 and 13 dpf larvae, respectively. **C.** Preference index (PI) across all eight directions for the two PNs shown in panel B. **D.** Radial plots showing developmental increase in CF direction selectivity index (DSI), at resultant angles calculated from the vector sum of PI. Red concentric rings mark the DSI from 0.2 to 1. Each line represents the DSI of a single PN. **E.** DSI distributions for three developmental groups. Median ± IQR : 0.245 ± 0.201 (early), 0.281 ± 0.322 (intermediate), 0.462 ± 0.305 (late) (Kruskal Wallis test for multiple comparisons (P = 0.003) followed by pairwise Mann-Whitney U test with Holm-Bonferroni correction for early vs intermediate (P = 0.363, Cohen’s d = 0.378), intermediate vs late (P = 0.089, Cohen’s d = 0.459), early vs late (P = 0.001, Cohen’s d = 0.96)). **F.** Left: Preference indices (normalized to cluster maximum) for three different CF inputs recorded from an example PN using whole-cell voltage-clamp recording. CF inputs recorded during the entire experiment were clustered based on the peak amplitudes. Right: Mean CF EPSC waveform for each cluster. Colours cyan, yellow and magenta represent clusters 1, 2 and 3 respectively. **G.** DSI values for all poly- and mono-innervated PNs obtained from whole-cell voltage-clamp recordings. For each cell, open black circle indicates the DSI calculated from all CF inputs, whereas filled circles indicate the DSI calculated for individual CF clusters. Fill color represents the mean CF EPSC amplitude of each cluster. **H.** DSI distributions for poly- and mono-innervated cells shown in panel G. Median ± IQR : 0.223 ± 0.282 (poly-innervated), (0.373 ± 0.225 (mono-innervated) (Mann-Whitney U test, P = 0.025, Cohen’s d = 1.529) N = 37, 37, 34 cells (from 19, 17, 17 larvae) for early, intermediate and late stages, respectively (B-E). N = 9,12 cells (from 6, 8 larvae) for early and late stages, respectively (F-H). Colours *green, grey* and *purple* represent early (4-5 dpf), intermediate (7-8 dpf) and late (11-14 dpf) larval stages, respectively. *P < 0.05, **P < 0.01, ns P > 0.05.

Next, we asked whether different fibers innervating a poly-innervated PN have distinct direction tuning profiles, and whether separating inputs from different fibers would change the tuning and resultant DSI. Whole-cell voltage-clamp recordings were performed from PNs at early and late stages while presenting optic flow in different directions as shown in Fig. 4A. To identify discrete CF inputs for each cell, we clustered the CF EPSC amplitudes using a gaussian mixture model. Cells with a single amplitude cluster were classified as mono-innervated, while those with more than one cluster were classified as poly-innervated (Fig. 4F-H). For all poly-innervated PNs, each cluster represented a unique fiber (Fig 4F, right). PI and DSI were calculated for each CF cluster (Fig 4F, left). We observed that indeed different CF clusters differ in their preferred direction/s (Fig 4F, left).

For each poly-innervated PN, we compared the DSI for all CFs received by the PN (overall DSI) to the DSIs of individual CF clusters (Fig. 4G). We found that clusters had distinct DSIs, with some higher and some lower than the overall DSI for the cell. As a result, overall DSI was always lower than the maximally tuned cluster (Fig. 4G). On the other hand, mono-innervated cells comprised of a single cluster whose DSI was identical to the cell’s overall DSI (Fig. 4G).

Next, we compared DSI of poly-innervated PNs with mono-innervated PNs (Fig. 4H). We found that mono-innervated PNs indeed have higher DSI (Fig. 4H). Consistent with the innervation patterns dominant at each developmental stage, the overall DSI of poly- and mono-innervated PNs mirrors that of early- and late-stage PNs, respectively (Fig. 4E, H).

These results indicate that when multiple fibers with different tuning properties converge on a single poly-innervated PN, it diminishes the PN’s direction selectivity. Consequently, once the PNs undergo pruning and become mono-innervated, their direction tuning improves as seen earlier in Fig. 4E. In sum, these results strongly suggest a causal relationship between CF pruning and the directional tuning of PNs.

### Simple plasticity rules can recapitulate pruning-driven sparsification and the emergence of directional selectivity in PNs

We propose that the developmental emergence of mono-innervation of PNs can arise through local activity-dependent plasticity under global resource constraints. To examine this, we simulated a bipartite network of N=20 inferior olivary units projecting onto N=20 PNs, with all-to-all initial connectivity. Synaptic weights evolved under a Hebbian term combined with subtractive normalization on both presynaptic and postsynaptic weight totals, capturing biological constraints of limited axonal/synaptic resources per olivary neuron and limited dendritic resources per PN. We drive developmental synaptic pruning through spontaneous, unstructured activity, modeled through weight updates under uniform random inputs to each CF.

We found that the network converged from dense, near-uniform connectivity to a sparse, near-bijective mapping in which each PN was dominated by a single CF (Fig. 5A, B). The simulated weight distributions transitioned from a poly-innervated to a mono-innervated regime over training, mirroring the developmental trajectory observed experimentally (Fig. 2D, E).

**Figure 5:**
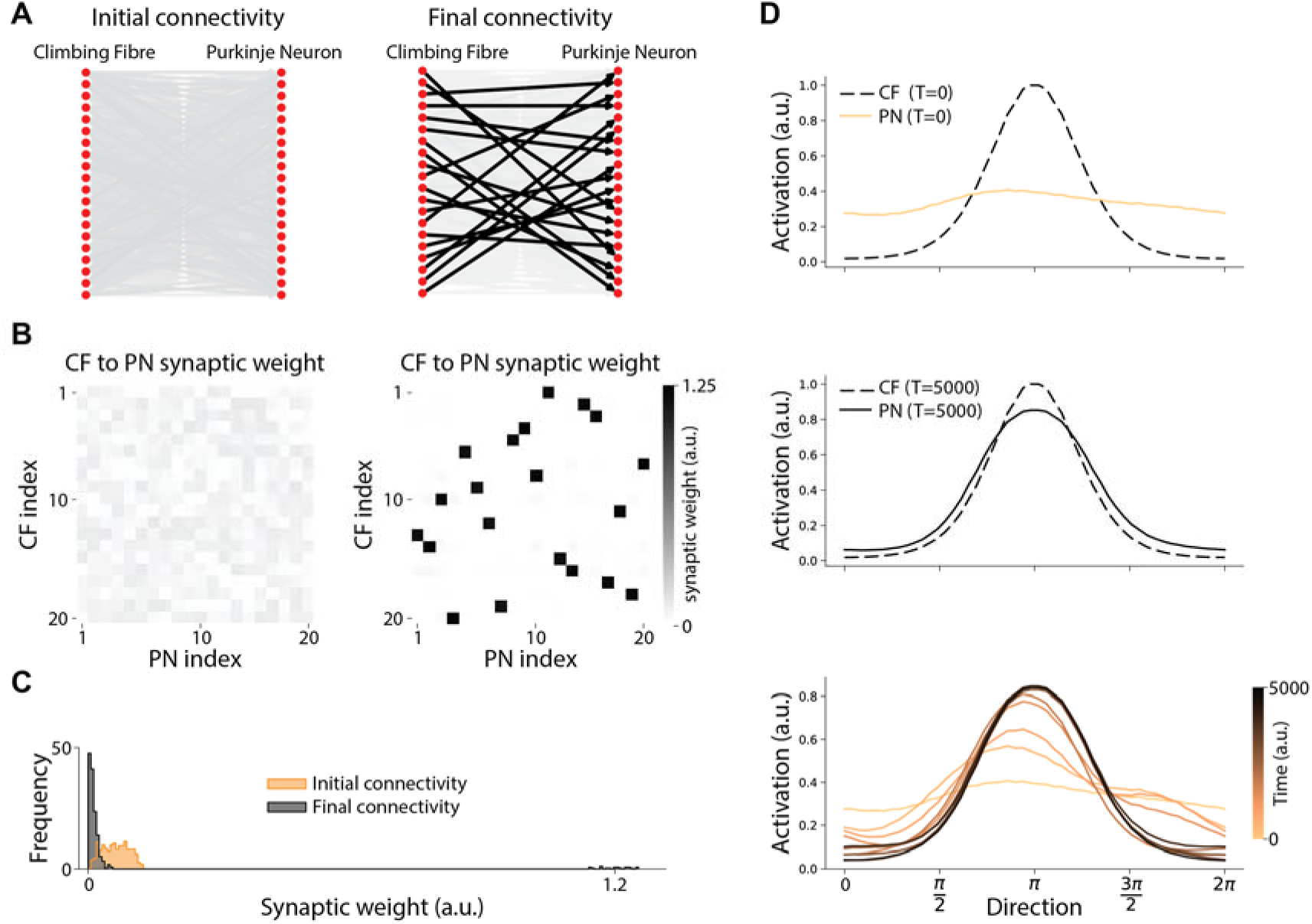
A minimal Hebbian model with pre- and post-synaptic resource constraints recapitulates pruning-driven sparsification and the emergence of directional selectivity in PNs. **A.** Bipartite network of *N* = 20 CFs and *N* = 20 PNs shown at initialization (left) and after *T* = 5000 iterations of developmental dynamics (right). Edge opacity is proportional to synaptic weight. Initial all-to-all connectivity refines to a near-bijective mapping in which each PN is dominated by a single CF. **B.** CF-to-PN synaptic weight matrix at initialization (left) and after training (right). Initial weights are weak and uniformly distributed; trained weights concentrate at one strong connection per PN column, with remaining entries near zero. **C.** Distribution of synaptic weights pooled across the matrix. The initial unimodal distribution near small values resolves into a bimodal distribution after training, with most weights eliminated and a small subset of dominant weights near 1.2. **D.** Tuning curves for an example CF (dashed) and its target PN (solid) probed with directional inputs. Top: at initialization, the PN integrates inputs from many directionally heterogeneous CFs and exhibits broad, weakly directional tuning despite the sharply tuned CF input. Middle: later in development, the PN inherits the directional tuning of its surviving dominant CF input. Bottom: the same PN’s tuning curve at successive training checkpoints (color, light to dark = early to late), showing progressive sharpening as pruning proceeds.

Correspondingly, the CF-to-PN synaptic weights evolved from an initially weak, near-uniform distribution to one with a small number of strong weights and the remainder near zero (Fig. 5C).

To assess how the synaptic refinement shapes sensory representations downstream, we then probed the network at successive training checkpoints with directional inputs in which each CF carried a von Mises-tuned response to a presented angle. Early in training, when each PN integrates inputs from many directionally heterogeneous CFs, PN tuning curves were broad and weakly directional. As pruning progressed, PNs progressively inherited the sharp directional tuning of their single dominant CF (Fig. 5D). Our model thus shows the emergence of directional selectivity in PNs, qualitatively consistent with the developmental increase in direction selectivity we observed experimentally (Fig. 4E).

Our model thus demonstrates that progressive mono-innervation and sharpened directional tuning are jointly producible by a single elementary learning rule operating under biologically plausible resource constraints — supporting the view that these functional outcomes need not arise from independent developmental programs but can emerge as joint consequences of the same competitive dynamics.

### Emergence of decorrelation in CF inputs to PN populations

Thus far, we have shown that representations of sensory input and sensori-motor error are refined during development at the level of single PNs. We next wanted to ask how these changes affect population level activity patterns. CF inputs cause large calcium transients that can be detected by calcium imaging (Fig. S4A) (*11*, *12*, *50*). Using single plane light sheet microscopy, we imaged spontaneous activity in GCaMP6s-expressing PNs at different developmental stages (Fig. S4B). Large calcium transients were used to estimate CF inputs (Fig. S4C). The spontaneous mean frequency of CF inputs estimated from calcium imaging (Fig. S4C, 0.255 ± 0.13 for early and 0.2 ± 0.111 for late stage, in Hz) was lower than what we measured from electrophysiology experiments (Fig. 1D, 0.4 ± 0.229 for early and 0.286 ± 0.216 for late stage, in Hz). Such an underestimation in frequency is expected due to slow kinetics of GCaMP6s (*51*).

Differences between preparations for electrophysiology and imaging may also contribute to the underestimation (see Methods). Nevertheless, consistent with electrophysiology, we saw a decline in the estimated frequencies over development (Fig. S4B, C). Late stage PNs exhibited significantly lower mean frequencies than the early stages (effect size = 0.602; comparable to effect size = 0.586 in Fig. 1D) (Fig. S4C). These results show that calcium imaging can be reliably used to resolve the changes in CF inputs to PNs caused by developmental pruning.

Pruning of surplus CF connections is expected to reduce shared inputs between different PNs, and, therefore desynchronize them and promote functional specialization. To assess the correlations in CF inputs among PNs, we computed Jaccard index (JI) between PN pairs from binarized time series of dF/F traces (see Methods). Consistent with previous studies in postnatal mice, JI decreased over development, suggesting a developmental desynchronization of PNs during the pruning window (Fig. S4D, Video S1) (*52*). Moreover, we observed that a significantly large number of PN pairs in the late larval stage were perfectly asynchronous (JI ≈ 0) (Fig. S4E) and the fraction of such pairs increased over development (Fig. S4F). Thus, developmentally, during the window of CF synaptic pruning, calcium transients in PNs become progressively more decorrelated. This may enhance computational efficiency by enabling distinct functions in PN clusters depending on the nature of signals carried by the winning CF. Such functional specialization is also evidenced from direction tuning (Fig. 4D, E).

### Synaptic pruning mediates efficient predictive processing

PNs receive directionally tuned sensory information and sensori-motor error responses from the olive which enables them to instruct motor learning, predictive processing and adaptive behaviors (*21*, *22*, *43*, *53*). We wondered if the developmental alterations in sensory tuning and sensori-motor error representations as shown above, will lead to behavioral consequences. We showed previously that when repetitive visual stimuli are presented, CF inputs to PNs ramp up in expectation of the arrival of that input. Similarly, when the expected input does not arrive, CF inputs encode an error in their expectation. Larvae use these predictive signals to learn stimulus patterns and generate swims with a short latency. When this circuit is compromised, larvae lose the ability to adjust swim latency (*22*). We tested how larvae at different developmental stages perform in this task and what the CF encoding of predictions and prediction errors looks like in PNs.

Towards this, we performed two-photon calcium imaging of PNs while larvae were engaged in a closed loop optomotor behavior with repetitive stimuli interleaved with probe stimuli (Fig. 6A) (see Materials and Methods). We first identified PNs that were forward or backward-tuned using a random optic flow stimulus set (Fig. S5A, B, see Methods). Next, we presented pulsatile strong forward optic flow (1cm/s) with eight repeats to acclimatize the larvae to this stimulus pattern. This was followed by three repeats of a ‘probe’ stimulus consisting of weak backward flow (-0.064cm/s) (Fig. 6B).

**Figure 6.**
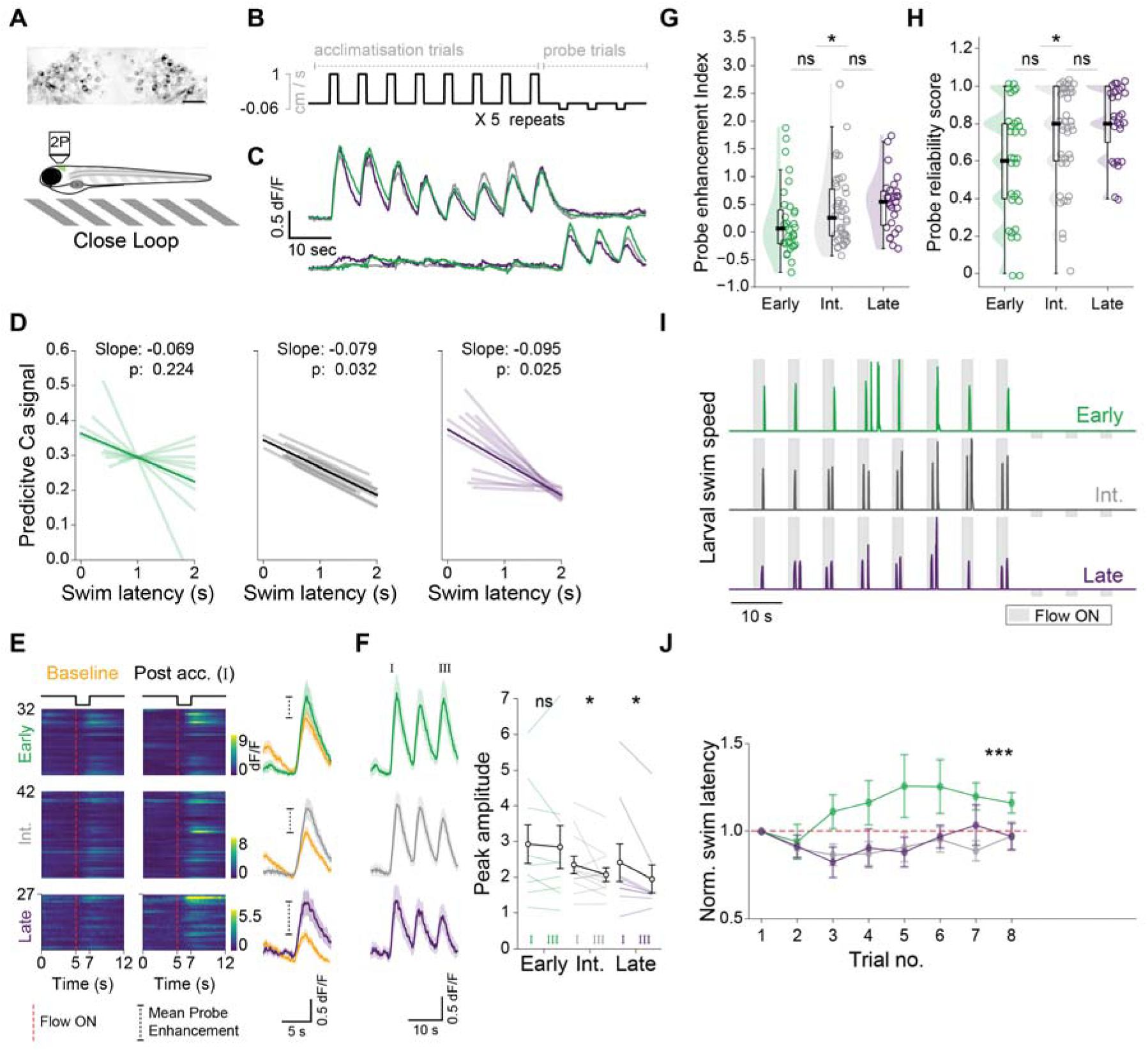
CF inputs to PNs shape sensory predictions and contribute to faster swim responses in late larval stages. **A.** Schematic of two-photon imaging in a closed-loop environment. Average GCaMP6s calcium activity in PNs of a transgenic aldoca:GCaMP6s larva imaged using two-photon microscope (top). Scale bar 20 µm. **B.** Optic flow trial structure. Five blocks of optic flow spaced by 30 - 40 s were presented to the larvae. Each block consisted of an acclimatisation phase (8 forward trials, speed 1 cm/s) followed by probe phase (3 backward trials, speed 0.064 cm/s). Each trial consisted of 5 s long stationary, followed by 2 s long moving gratings. **C.** Average dF/F traces from forward (top) and backward (bottom) tuned PNs during acclimatisation and probe trials at different larval stages. **D.** Emergence of small but significant negative correlation between swim latency and pre-flow calcium activity in forward tuned PNs over development. Mixed-effects linear models with random intercepts computed for N = 10, 10, 12 larvae for early, intermediate and late larval stages, respectively. **E.** Enhanced probe response in backward tuned PNs over development. Left: Heat map showing average dF/F for backward tuned PNs over development. Probe enhancement index was calculated as fractional change in peak dF/F during the first probe trial after acclimatisation over a baseline (orange, calculated from 5 random probe trials shown in Fig S5). Right: dF/F traces for baseline and first probe trial (Mean ± SEM). Red dashed line indicates the start of the first probe trial. **F.** Decay in probe response over three probe trials. Left: dF/F traces for backward tuned PNs (Mean ± SEM). Right: Comparison of peak amplitude in first (I) and third (III) probe trials. One-sided Wilcoxon signed-rank test was performed to test the probe decay for early (P = 0.285, Cohen’s d = 0.047), intermediate (P = 0.046, Cohen’s d = 0.36) and late (P = 0.013, Cohen’s d = 0.333) larval stages. N = 9, 12, 9 larvae from early, intermediate and late stages, respectively. **G.** Distribution of probe enhancement indices of backward tuned PNs over development. Median ± IQR : 0.063 ± 0.61 (early), 0.256 ± 0.839 (intermediate) , 0.549 ± 0.62 (late) (Kruskal Wallis test for multiple comparisons (P = 0.026) followed by pairwise Mann-Whitney U test with Holm-Bonferroni correction for early vs intermediate (P = 0.234, Cohen’s d = 0.309), intermediate vs late (P = 0.234, Cohen’s d = 0.212), early vs late (P = 0.027, Cohen’s d = 0.567)). **H.** Increase in probe reliability score (fraction of blocks where probe response was higher than baseline) of backward tuned PNs over development. Median ± IQR : 0.6 ± 0.4 (early), 0.8 ± 0.4 (intermediate), 0.8 ± 0.3 (late) (Kruskal-Wallis test for multiple comparisons (P = 0.016) followed by pairwise Mann-Whitney U test with Holm-Bonferroni correction for early vs intermediate (P = 0.058, Cohen’s d = 0.525), intermediate vs late (P = 0.522, Cohen’s d = 0.256), early vs late (P = 0.021, Cohen’s d = 0.811)). N = 32, 42, 27 backward tuned cells (from 10, 12, 13 larvae) for early, intermediate and late stages, respectively. **I.** Normalised swim speed (between 0 to 1) of early, intermediate and late stage larvae in response to a single block of optic flow. **J.** Average swim latency normalized to the first trial of the acclimatisation phase. Red dashed line indicates no change from the first trial. N = 10, 12, 13 larvae for early, intermediate and late larval stages, respectively. Linear mixed-effects model with ANOVA, including trial number and trial number plus larval age showed a significant effect of age on swim latency over trials (P < 0.001). Colours *green, grey* and *purple* represent early (4-5 dpf), intermediate (7-8 dpf) and late (11-14 dpf) larval stages, respectively. ***P < 0.001, **P < 0.01, *P < 0.05, ns P > 0.05.

The forward and backward tuned PNs identified earlier (Fig. S5B) showed a rise in average calcium activity during the acclimatization and probe phase, respectively (Fig. 6C). In line with direction selectivity, these calcium transients have been previously attributed to climbing fiber inputs in similar experiments (*22*). We observed these calcium transients in all three developmental stages (Fig. 6C). The calcium signal in forward-tuned PNs prior to forward-flow onset is predictive of swim latency in intermediate and late-stage larvae (Fig. 6D). That is, higher the calcium signal in PNs prior to forward flow onset, lower the swim latency. However, such a negative correlation is absent in early-stage larvae indicating that PNs poly-innervated by CFs lack the ability to encode the swim latency predictively (Fig. 6D). Second, acclimatization to forward flow enhances the response to backward flow during the probe phase in backward-tuned PNs (Fig. 6E) and, this enhanced response decays over the probe phase (Fig. 6F). We defined baseline response as the response of backward tuned PNs to weak backward flow during the presentation of random flow stimuli in the first block (Fig. S5). Compared to baseline, the response to the first probe stimulus post acclimatization was significantly enhanced in intermediate- and late-stage larvae (Fig. 6E). Such enhancement was absent in early-stage larvae (Fig. 6E, G). The extent of probe enhancement gradually increased, with late-stage PNs showing significantly larger values (effect size = 0.567) compared to the early stage (Fig. 6G). Similarly, the reliability with which probe enhancement occurred across trials was also increased in late-stage larvae compared to early-stage larvae (Fig. 6H). In sum, it appears that PNs at stages where they are poly-innervated by CFs are poorly equipped to acquire models of sensory stimuli. They are also poor at detecting errors in stimulus patterns when expectation does not match reality. In our earlier study, we showed that the ability to predict an upcoming sensory stimulus leads to quicker swim responses. Larvae respond with swims to forward optic flow stimuli but not to the probe stimuli (Fig. 6I). With repetitive forward flow stimuli, intermediate- and late-stage larvae, but not early-stage larvae, were able to decrease their swim latency (Fig. 6J). Taken together, our results show that CF inputs arising from multiple fibers in poly-innervated PNs convey poorly tuned sensori-motor information, resulting in an inability to modulate swim latency. The single CF inputs in mono-innervated PNs of late stage larvae are able to deliver specific sensori-motor error information to aid in motor planning, leading to faster behavioral responses.

## Discussion

### Selective pruning improves signal fidelity

By examining individual CF inputs within poly-innervated PNs, we found that convergence of differently tuned inputs degrades the fidelity of downstream sensory and motor representations. As CF inputs to PNs are pruned away, convergence of differently tuned CFs is reduced, the fidelity and temporal precision of sensorimotor signals are improved, providing a circuit substrate for the emergence of predictive representations. These changes may enhance the ability of the olivo-cerebellar circuit to form predictive representations of regularities in the sensory environment, thereby supporting motor planning.

Developmental synaptic pruning is widespread-it is known to occur at the neuromuscular junction, in the autonomic nervous system, in the lateral geniculate nucleus and in the neocortex, apart from the cerebellum (*1–3*, *6*, *7*, *9*) and often reduces convergence and shared connectivity. Our findings suggest one possible functional consequence of this process: reducing convergence among differently tuned inputs can increase the selectivity and temporal precision of neuronal representations, making such neural representations more behaviorally relevant.

### Advantages of the zebrafish model

The zebrafish model used in this study enabled us to examine the functional consequences of synaptic pruning in vivo during a period when the developing animal is already engaged in active sensorimotor behavior. In zebrafish, Purkinje neurons are specified by 2.5 to 3 dpf and functional CF inputs can be recorded by 4 dpf (*28*, *29*). A layered cerebellar structure appears, new PNs continue to be born and PNs elaborate their dendritic arbors by 5 dpf. PN neurogenesis and growth of arbors continues at least until 8 dpf (*32*, *54*). During this time, PNs are electrically active and participate in many motor behaviors such as OMR and optokinetic response (*22*, *29*, *43*, *55*). Thus, in larval zebrafish, the cerebellum is functional, yet undergoing critical developmental modifications during these early larval stages.

We show here that in zebrafish, PNs proceed from poly-innervation by CFs to mono-innervation during the first two weeks of life. This is consistent with previous observations in zebrafish (*29*, *33*) and in line with a similar developmental window in mammals (*14*). We also find a small proportion of PNs with more than one CF in the late-larval stage. It is worth noting that a recent study showed that some adult mouse PNs and a large proportion of adult human PNs are innervated by two CFs on two primary dendritic branches of a single PN (*58*). It would be interesting to investigate whether adult zebrafish PNs are indeed innervated by one or two CFs.

### CF synaptic pruning is one among many developmental changes in the cerebellum

The cerebellum undergoes several major changes during the first two weeks of life as is known from studies in rodents and zebrafish. In the rodent cerebellum, transient synaptic connections between PNs can be seen in the first post-natal week, which are reduced in number in the second post-natal week and entirely absent in the adult. Such connections have been suggested to be important for the propagation of traveling waves of activity (*56*, *57*). In addition, there are many other metamorphic events such as conversion of GABA from being depolarizing to inhibiting, the neurogenesis and migration of granule cells, Purkinje neuron dendritic elaboration, etc., (*56*) In zebrafish, as noted above, the period of CF to PN synaptic pruning coincides with periods of PN neurogenesis, PN dendritic growth, neuromodulatory innervation, etc., All of these changes may contribute to progressive evolution of cerebellar computation, necessitating further in depth studies of these phenomena.

## Conclusion

Together, our findings suggest that developmental synaptic pruning can do more than reduce excess connectivity: by selectively eliminating competing inputs, it can sharpen neuronal representations and help establish the computational properties of mature circuits. The emergence of predictive sensorimotor signals during this refinement provides an example of how developmental changes in connectivity may enable increasingly sophisticated neural computations and adaptive behavior.

## Materials and Methods

### Biosafety and Ethics

All experiments in this study were performed on larval zebrafish (*Danio rerio*) from 4-19 dpf with approval of the Institutional Animal Ethics Committee (NCBS-IAE-2020/14(R1E)) and Institutional Biosafety Committee (TFR/NCBS/14-IBSC/VT1/2011). ZFIN was used as a reference for all zebrafish protocols (zfin.org). The following fish lines were used for different experiments as appropriate: hspGFFDMC28C:gal4; UAS:gfp (Prof. Koichi Kawakami, National Institute of Genetics, Japan) (*62*); aldoca:GCaMP6s; nacre^-/-^ (Prof. Masahiko Hibi, Nagoya University, Japan); ca8-cfos construct for driving expression in Purkinje neurons (Prof. Hideaki Matsui, Niigata University, Japan) (*55*).

### Animal Maintenance

Adult zebrafish were housed in ZebTec zebrafish housing system (Tecniplast, Italy) at a temperature of 28°C, pH 7.4, conductivity of 1200 µS and 14:10 hrs light-dark cycle. All adult fish were fed twice a day with freshly hatched brine shrimp (Summit Artemia, Great Salt Lake Artemia, USA) in the morning and adult zebrafish diet (Zeigler, USA) in the evening. Evening feed was replaced with freeze dried bloodworms, (Hallofeed, India) twice a week. Juvenile fish were given the same morning diet plus three feedings of larval AP100 <100 microns (Zeigler, USA) interspersed during the day.

Embryos were rinsed thoroughly and raised in E3 medium treated with 0.36 µM methylene blue (M9140, Sigma-Aldrich, USA) in 90mm petri dishes (Tarsons Products, India). Up to 15 days post fertilization (dpf), larvae were housed in a 28°C incubator (MIR-154, Sanyo, Japan) with a zeitgeber cycle identical to adults. Developmental synchronization was performed at three stages- (a) 24 hours post fertilization, 5 - 26+ somite stage, (b) 2.5 - 3 dpf, hatched larvae, and (c) 4 dpf, larvae with a visible swim bladder. At this stage, the larvae were transferred to tanks (12 x 8 x 5 cm) for experiments and thereafter, fed with larval AP100 <100 microns four times a day. Population of 25-35 larvae per tank was maintained at all times and E3 medium was replaced every day.

### Developmental stages

Three larval stages were used in this study, namely, 4-5 dpf (early larval stage), 7-8 dpf (intermediate larval stage) and 11-14 dpf (late larval stage; up to 19 dpf for fictive swim recordings).

### Visualization of CFs and PNs

The inferior olivary nucleus was labelled with green fluorescent protein (GFP) in a transgenic hspGFFDMC28C:gal4; UAS:gfp larva. Sparse genetic labelling of olivary neurons was achieved by microinjecting single-celled embryos with Tol2-UAS:mcherry plasmid (50 ng/μL) and Tol2 transposase mRNA (100 ng/μL) (*63*). PNs were labelled using Tol2-ca8-cFos:ecfp/dendra2 plasmid (50 ng/μL for sparse genetic labelling). Imaging was performed using light-sheet (Zeiss Lightsheet 7) and confocal (FV3000, Olympus, Japan) microscopes. Acquired images were merged in Fiji (*64*).

### Preparation for electrophysiology

Larval zebrafish were anaesthetized using 0.0168% w/v tricaine/ MS-222 (E10521, Sigma-Aldrich, USA). Micro-dissections were performed on live anaesthetized fish under a stereo microscope (SZX16, Olympus, Japan) with oblique illumination. Using short tungsten pins (M249400, California Fine Wire Company, USA, 25.4 μm), fish were positioned laterally on a sylgard (SYLGARD^TM^ 184 Silicone Elastomer Kit, The Dow Chemical Company, Germany) base in the center of an in-house custom designed acrylic recording chamber or a transparent 60 mm petri dish (Corning, USA). Two pins were first pushed through the notochord, one at a rostral segment adjacent to the swim bladder and another at a caudal segment. As per the requirement of the preparation (dorsal-up/ ventral-up, described below), the head was rotated, tricaine was washed out with external saline (composition in mM; 134 NaCl, 2.9 KCl, 1.2 MgCl2, 10 HEPES, 10 Glucose, 2.1 CaCl2; 290 mOsm, pH 7.8). The pinned larva was then paralyzed by adding a drop of 1 mg/mL α-bungarotoxin (203980, Sigma-Aldrich, USA) dissolved in external saline for 7-10 minutes following which it was washed with external saline. Alternatively, 0.016 mg/mL tubocurarine (T2379, Sigma-Aldrich, USA) was used in the external saline throughout the recording. In the dorsal-up preparation, skin from the dorsal side of the brain was peeled using fine forceps (Student Dumont #5, Fine Science Tools, Canada) to expose the cerebellum. In the ventral-up reduced preparation, ventral tissue was gently removed with the help of a tungsten pin and fine forceps just until the brain was accessible. The brain, eyes, and the trunk were kept intact during this procedure (adapted from (*65*)). The recording chamber containing the larval preparation was placed on a fixed stage (Scientifica, UK) for single cell and fictive motor recordings.

### Single cell recordings

Patch pipettes were made by pulling filamented borosilicate glass (1.5 mm OD, 0.86 mm ID; Harvard Apparatus, USA) using a Flaming/ Brown P97/P1000 puller (Sutter Instrument, USA). Pipettes were backfilled with either of two internal salines to achieve a pipette resistance of ∼10 MΩ: cesium gluconate internal saline (composition in mM: 115 Cs hydroxide, 115 Gluconic acid, 15 CsCl, 2 MgCl2, 10 HEPES, 10 EGTA, 4 MgATP; 290 mOsm, pH 7.2) was used for voltage-clamp recordings in graded stimulation experiments. For current-clamp and voltage-clamp recordings in open-loop motor experiment, we used K-gluconate internal saline (composition in mM: 115 K gluconate, 15 KCl, 2 MgCl2, 10 HEPES, 10 EGTA, 4 MgATP; 290 mOsm, pH 7.2). Sulforhodamine (S1402, Sigma-Aldrich, USA) at final concentration of 5 μg/mL was added to the internal saline and filtered using 0.22 μm filter.

The recording pipette was positioned using a motorized micromanipulator (Double PatchStar, Scientifica, UK). Neurons were approached with the help of a 60x water immersion objective (CFI Fluor 60X, 1.0 NA, Nikon, Japan) on a Nikon microscope (Eclipse FN1 Upright, Nikon, Japan) connected to a Flea3-USB3.0 camera (FLIR, USA).

In voltage clamp experiments, the holding potential was set to -55mV. Passive properties (series resistance, input resistance and input capacitance) were calculated from a 150-200 ms long hyperpolarizing pulse to -80 mV delivered intermittently during the experiment. For all recordings in this study, series resistance was less than 20% of input resistance. Over the course of the experiment, not more than 25% change in series and input resistances was allowed. In addition, recordings were monitored over time to ensure stable CF EPSC amplitudes. Recordings of spontaneous CF EPSCs were performed in the dark for 45 to 180 s. Loose patch recordings were performed as above but with the pipette filled with external saline. Upon establishing a stable seal of 60-100 MΩ, recordings were performed in I=0 mode.

### Fictive motor recordings

Suction pipettes were made from non-filamented thin-walled glass (1.5 mm OD, 1.10 mm ID; Harvard Apparatus, USA) pulled to a tip diameter of 30-40 μm. The pipette was backfilled with external saline and positioned at the inter segmental junction with the help of the camera. Gentle suction was applied and locked to get stable recording from the ventral root of the spinal nerve. All suction recordings were obtained from ± 2 segments from the cloaca. Spikes with an inter spike interval (ISI) of 1 s were classified into different swim bouts. Bouts with at least 3 spikes were considered for analysis.

### Data acquisition and analysis

All electrophysiology data were acquired using Multiclamp 700B amplifier, Axon Digidata 1440A and Axon pCLAMP 10 acquisition software (Molecular Devices, USA) at a sampling rate of 20 or 50 kHz with a low pass filter of 1.8 or 2 kHz. The gain was set to 1 for whole-cells, 200 for loose-patch and 200 - 2000 for fictive swim recordings. Acquired data was stored as .abf files and analyzed post hoc with custom Python scripts. Spike detection for loose-patch recordings was performed using MATLAB functions from the open access mbl-nsb-toolbox (https://github.com/wagenadl/mbl-nsb-toolbox).

### Graded CF stimulation

Climbing fibers were electrically stimulated in a dorsal up larval preparation with the help of a twisted pair bipolar electrode custom-made using Formvar insulated Nichrome wire (76100, A-M Systems, diameter-17.78um bare, 25.4um coated). The bipolar electrode was positioned at the lateral tip of the cerebellar lobe and simultaneous whole-cell recording was performed from a single ipsilateral PN (Fig. 2B).

CFs were stimulated at a range of current intensities to create electric fields of varying strengths. Depending on the strength of the electric field and relative distance of a fiber from the center of the field, single action potentials were stochastically elicited and a CF EPSC was recorded in the PN with a latency of 2-5 ms. The number of discrete steps of evoked CF EPSC amplitudes seen in PNs was used as a readout of the innervation. For each current intensity, ten pulses, 100 µs long, were delivered at a frequency of 0.2 Hz by a stimulus isolator (ISOflex, A.M.P. Instruments, Israel). Maximum and minimum intensities were identified as values at which all ten trials resulted in either the largest possible evoked CF EPSC amplitude (100% success) or no evoked CF EPSC at all (100% failure), respectively. All current intensities between maximum and minimum values were sampled with 10 µA steps. Sequence of intensities was randomized to prevent effects of short-term plasticity. The current intensities ranged between 10 - 480 µA. Trials with noisy baseline, overlapping EPSCs, or large stimulus artefacts were excluded from analysis.

### Anatomical evidence for poly-innervated PNs in larval zebrafish cerebellum

Previously reported correlated light and electron microscopy (CLEM) dataset was used to extract examples of poly-innervated PNs (*34*). PNs with characteristic dendritic arbors were identified and then their pre-synaptic CF partners were searched using the annotated synapses. Segment IDs for example neurons are as follows: Fig. 2A (left): 7205847396 (PN), 8674447299, 7185454674, 8715310474 (CFs). Fig2A (right): 7287445731 (PN), 10265648783,10265654529, 13366327969 (CFs).

### Innervation index and amplitude disparity index

The innervation index was computed as the sum of CF steps normalized to the maximum step amplitude. This results in an innervation index of 1 for mono-innervated PNs. Values close to 1 suggest poly-innervation dominated by a strong CF alongside weaker CFs, while substantial deviations from 1 indicate poly-innervation with multiple strong CFs.

Amplitude disparity index was computed to measure the fold change between the strongest CF EPSC and the weaker ones as follows:

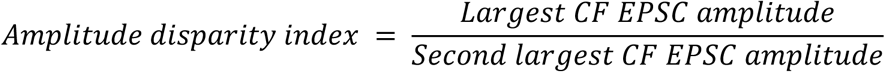

A large disparity index indicates selective strengthening or maintenance of a single CF while weakening the others.

### Paired-pulse experiment

Paired pulse experiment was performed by stimulating climbing fibers in a similar configuration as graded stimulation experiment. Initially, maximum intensity (current value that results in 100% success) was identified by delivering five 0.1 ms long pulses at a frequency of 0.2 Hz.

Subsequently, maximum intensity trials, each consisting of two pulses separated by an interstimulus interval (ISI) were delivered. Trial frequency of 0.1 Hz and ISIs of 30, 35 and 50 ms were used. In addition to the trial exclusion criteria in the graded stimulation experiment, trials where one or more spontaneous CF EPSCs occurred in the interstimulus interval were also excluded. Paired Pulse Ratio (PPR) was calculated from the EPSC amplitudes evoked by the two current pulses (*I* & *II*) as follows:

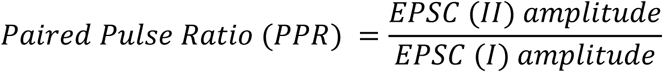

For each ISI, we measured the mean PPR from 3-10 trials.

### Optic flow presentation

Optic flow consisted of moving black and white gratings with a spatial period of 1 cm on a screen. A custom Processing (www.processing.org) script was used to create the gratings and update their position at a frequency of 120 Hz (which is also the refresh rate of the screen). For experiments with electrophysiology, a 5-inch Raspberry Pi LCD display was used to present whole field optic flow beneath the larva with a pixel size of 0.13 mm. For experiments with two-photon imaging, black and red gratings were projected on a diffusive screen with a pixel size of 0.32 mm, using a modified LCD screen (as described previously in (*22*)). Each trial of optic flow comprised stationary followed by moving gratings. Duration and direction of moving and stationary phases are indicated in the description of the respective experiments.

### Open-loop optic flow presentation with electrophysiology

Fictive motor recordings were obtained from dorsal up-preparations as described above. The objective was then gently moved along the body axis to focus on the cerebellum. Upon securing a stable recording from a single PN, the LCD display mounted on a micromanipulator (MM-3, Narishige, Japan) was gently slid under the recording chamber containing the larval preparation. For the experiment in Figure 3, optic flow consisted of seven trials of forward moving gratings in an open-loop environment. Each trial was 20 s long, with a 10 s stationary followed by 10 s moving gratings. Simultaneous suction and whole-cell recording with visual stimulus were performed in both voltage- and current-clamp mode. CF frequencies were calculated from both voltage- and current-clamp recordings, while CF amplitude analysis was performed only on voltage-clamp data.

For the experiment in Figure 4, directional optic flow was presented in an open-loop environment to the larvae in dorsal up preparation. The experiment consisted of five blocks of visual stimulus. Each block consisted of eight randomized trials in one of the eight directions, i.e., forward, backwards, left, right, forward left, forward right, backwards left and backwards right. As a result, five trials were presented for each direction. Each trial consisted of a 5 s stationary period followed by a 3 s flow period. Direction was changed at the onset of the stationary phase in each trial. Simultaneous loose-patch or whole-cell voltage-clamp recordings were performed from PNs alongside presentation of directional stimulus.

### Motor Tuning Index

Motor Tuning Index (Fig. 3G, H) was defined as the difference between the CF frequency in the motor window and the mean of the pre- and post-motor CF frequencies, normalized by their sum as described under:

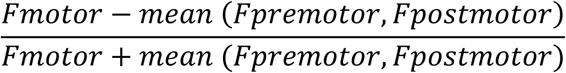

Here, F_premotor_, F_motor_, F_postmotor_ are the CF frequencies during the long pre-motor, motor and post-motor windows, respectively-each 300 ms long.

### Direction Selectivity Index (DSI)

CF frequency was calculated for each trial in the 1.5 s following flow start. This period is expected to capture sensory associated CF inputs (Fig. 4B). For each direction, Preference Index (PI) was calculated as the product of the mean CF frequency across 5 trials and the fraction of trials with non-zero CF frequency. Direction Selectivity Index (DSI) was computed from PI in vector space:

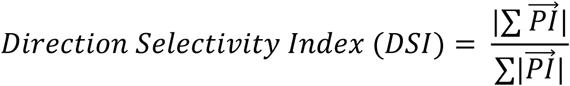

DSIs range between 0 and 1, where 0 represents a neuron with equal response in all eight directions and 1 represents a neuron that responds only in a single direction (*19*, *66*).

### Computational model of CF–PN refinement

We simulated a bipartite network of *N*_CF_ = *N*_PN_ = 20units with all-to-all weighted connectivity 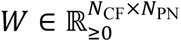, initialized as *W*_ij_ ∼ U(0,0.1) + 10^−3^.

On each iteration, CF activity *x* ∈ ℝ^NCF^ was drawn as *x*_i_ ∼ U(0,1) i.i.d., representing unstructured spontaneous activity. PN activity was computed as *y* = tanh(*W*^T^*x*). Weights were updated according to a Hebbian rule with subtractive normalization on row and column sums:

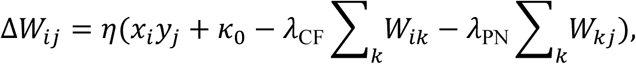

with *η* = 0.05, n_0_ = 1, *λ*_CF_ = *λ*_PN_ = 0.5. Weights were clipped to be non-negative after each update. The presynaptic (*λ*_CF_) and postsynaptic (*λ*_PN_) normalization terms implement, respectively, a per-CF axonal resource constraint and a per-PN dendritic territory constraint. This plasticity mechanism was driven for *T* = 5000 iterations for independently drawn random CF activity.

To assess how the evolving connectivity would shape sensory representations, we assigned each CF to have a preferred angle *θ*_i_ = 2*πi*/*N*_CF_. Then, the network was probed at various time points with directional inputs provided to the CFs. For a directional input angle ф, each CF *i* produced a response of *x*_i_ ∝ exp[ncos(*θ*_i_ — ф)] with n = 2, normalized by the population maximum.

Tuning curves for individual PNs were obtained by presenting 50 probe stimuli spanning [0,2*π*)and recording *y*_j_ = tanh((*W*^T^*x*)_j_). These probe inputs were not used to update weights; they served only to read out the directional selectivity inherited by each PN from the surviving CF connections.

### Agarose embedding

Transgenic aldoca:GCaMP6s; nacre^-/-^ larvae were first anaesthetized and then paralyzed using using 0.0168% w/v tricaine/ MS-222. For light-sheet imaging, larvae were paralyzed using 1 mg/mL α-bungarotoxin and then embedded vertically inside a thin glass capillary using 1% low gelling agarose (A9414, Sigma-Aldrich, USA) made in E3 medium. The capillary was then held in the sample holder of a light-sheet microscope (Lightsheet 7, Zeiss, Germany) and the fish was lowered in a chamber filled with filtered system water and the cerebellum was located.

For two photon imaging, a drop of 2% low gelling agarose made in E3 medium was used to embed the larvae in a transparent 60mm petri dish (Corning, USA) facing dorsal side up. After 20 minutes, the dish was filled with E3 medium. The tail was then freed by carefully removing the agarose around the swim bladder using a surgical blade. The embedded larvae were allowed to acclimatize for 7-10 hours before the experiment.

### Light-sheet calcium imaging

Spontaneous activity was imaged for 1.5 minutes in PNs. A single plane was imaged using pivot scan mode with 4-10% of the input laser power (25mW) on the Zeiss Zen Black acquisition software. Dual side (left and right) illumination was performed using a pair of 10x air objectives (LSFM 10x/0.2 foc, Carl Zeiss, USA). Emission was detected orthogonally using a 20x water immersion objective (W Plan-Apochromat, 1.0 NA, Carl Zeiss, USA). Online fusion was carried out from the images captured by left and right illumination. The resulting sampling rate was 25 Hz.

### Two-photon imaging of PNs

GCaMP6s was excited using a Titanium:Sapphire laser (Chameleon Ultra II, Coherent Inc, USA), tuned to 920 nm with a peak pulse width of 140 femtoseconds, frequency of 80 MHz and power of up to 30mW at sample. ScanImage 3.8 (*67*) was used to perform time series imaging of a single plane of PNs. A field of view with 512*128 pixels was scanned using pixel size of 0.32 µm at a frame rate of 7.81 Hz (resultant pixel dwell time of 1.6 µs). Image acquisition was synchronized with behaviour experiments using a custom Processing script that sends a TTL pulse to ScanImage. In order to acclimatize the larvae to high frequency vibration of XY-scan mirrors, optomotor behaviour experiments were programmed 30 s after the initiation of two-photon imaging. Imaging was performed continuously for up to 15 minutes.

### Optomotor behavior

For behaviour experiments performed along with two-photon imaging, a small square shaped window was cut on the projection screen underneath the larva. This allowed high speed video recording of the tail at 200 frames per second (fps) using a Flea3-USB3.0 camera (FLIR, USA). With the help of Bonsai (*68*), consecutive frames in the video recording were subtracted and processed to get real-time Larval Locomotor Drive (LLD). Open Sound Control (OSC) communication was then used to send LLD to the Processing script, running in parallel. In closed-loop (CL) environments, this Processing script compensates for the larval swims when updating the gratings, while in open-loop (OL) environments, the updates are independent of larval swims.

### Estimation of larval swim velocity

During CL behaviour experiments, larvae are allowed to engage with the optic flow. This was done by compensating for the grating movement based on larval swim velocity. To estimate the swim velocity, LLD was scaled to an average velocity of 1.5 cm/s. Scaling factor was calculated by presenting a 10 s long forward optic flow in OL environment to each larva before beginning of close loop experiment, as:

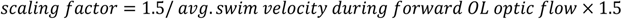

Using this scaling factor, raw-Instantaneous Swim Velocity (ISV_raw_) was computed for each frame *n*, as follows:

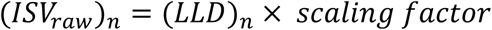

Thereafter, a rolling exponential smoothing algorithm was implemented to get Instantaneous Swim Velocity (*ISV_n_*) for each frame.

### Closed loop feedback

Optic flow presentation was structured uniquely by experimentally setting the flow, gain, grating speed, grating direction. For a given frame *n*, the grating displacement (d_grat_)_n_ was computed as:

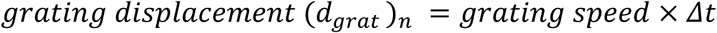

Here, *Δt* represents the time interval between previous and current frame (equals roughly 8 ms, resulting from iterations at 120 Hz in Processing). ISV_n-1_ was initialized to 0 for the first frame at the start of the experiment.

Displacement of the larva was estimated as:

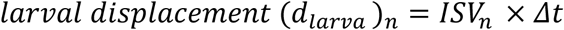

The updated grating position (Pos_n_) was computed by compensating for the larval displacement as follows:

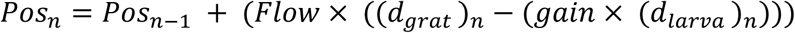

Gain was set to 0 for open loop and 1 for closed loop. Flow was set to 0 for stationary gratings and 1 for moving gratings.

### Prediction-error experiment

The experiment was performed in two blocks separated by 30 s. First block consisted of five random repeats for each of the following trial types: forward flow with grating speed of 1 cm/s, forward flow with grating speed of 0.064 cm/s, backward flow with grating speed of 1 cm/s and backward flow with grating speed of 0.064 cm/s. This block was used for classification of cells as forward, backward or motor (Fig. S5A, B). The second block comprised five repeats, each consisting of an acclimatization phase followed by a probe phase (Fig. 6B). During the acclimatization phase, eight forward trials were presented with a grating speed of 1 cm/s. During the probe phase, three backward trials were presented with a grating speed of 0.064 cm/s. The five repeats were separated by an interval of 30-40 s. All trials had 5 s stationary followed by 2 s moving gratings (*22*).

### Extraction and analysis of GCaMP signals

Time series of images acquired from the light-sheet and two-photon microscope were pre-processed in Suite2p (*69*) by optimizing the parameters for cell detection (Table S1). Motion correction was performed using non-rigid registration in blocks of 128 x 128 (light-sheet) or 64 x 64 (two-photon) pixels. Cellpose (*70*) anatomical detection method was used to find masks on enhanced mean images. ROIs were detected with a diameter ∼6 microns and those with no overlapping pixels were only used for analysis. Semi-automatic identification of cell bodies was performed using Suite2p GUI. Mean neuropil fluorescence (Fneu) was estimated using inner neuropil radius equal to soma radii. For each ROI, 0.7 times Fneu was subtracted from mean fluorescence value to get frame-wise ROI fluorescence (F) which was then NaN-padded on both ends. Baseline was calculated as the tenth percentile of ROI fluorescence (F) in a 1-minute window around each frame. Fold change in fluorescence from baseline (dF/F) was computed framewise for each ROI as follows:

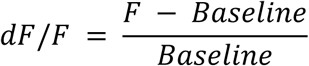

dF/F trace was smoothened using a Gaussian filter (SciPy ndimage gaussian_filter1d, sigma=3) for 1P imaging (*71*).

### Calculation of Jaccard index

1.5 minutes long time series light-sheet images were used to compute synchrony between PNs using Jaccard Index. To ensure a good signal-to-noise ratio, cells with at least 20% dF/F were selected. dF/F peaks were detected with the help of the find_peaks function in SciPy (*71*). Peaks were assigned as CF associated calcium transients using a threshold and prominence of 5%. For active cells with 3 or more peaks, the dF/F for all frames from peak minus 200 ms to peak were set to one and the other frames were set to zero, resulting in a binary activity time series. The degree of overlap between a pair of neurons A & B was computed using Jaccard index (intersection over union) with the help of the Scikit-learn Python module (*72*, *73*).

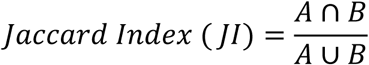

Jaccard index was calculated for all possible pairs of active PNs within a cerebellar lobe for each larva. Pairs with JI < 0.01 were assigned as asynchronous.

### Classification of PNs as motor, forward or backward tuned

PNs were classified as motor, forward or backward tuned from the first block of stimulus presentations in random order (Fig. S5). Forward and backward regressors were generated by convolving the respective components of the optic flow with the GCaMP kernel, which was modelled as a dual alpha exponential function (Fig. S5A). Similarly, a motor regressor was generated for each larva using the swim velocity. Thereafter, multiple linear regression was performed to fit the calcium activity of individual PNs to the motor, forward and backward regressors. PNs with R2 0.4 were classified as motor-, forward- or backward-tuned based on the largest regression coefficient (Fig. S5B) (*22*, *50*).

### Calculation of probe enhancement and reliability score

The Probe Enhancement (PE) index was calculated for all backward tuned PNs using the peak dF/F values from before and after the acclimatization, as given below:

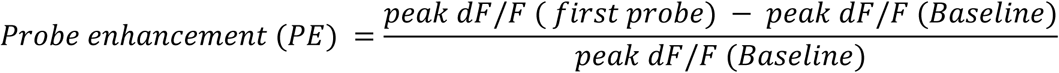

The peak dF/F (Baseline) was calculated as the average response across five backward trials presented at 0.064 cm/s during the first block of the prediction-error experiment (Fig. S5). The peak dF/F (first probe) was calculated as the average response across five trials of first probe trial during the second block of the experiment (Fig. 6B).

In addition, to assess the fidelity with which probe enhancement was seen upon acclimatization, we computed a reliability score over the five repeats as:

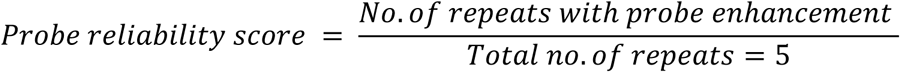

### Statistical analysis

Linear mixed-effects model with ANOVA (Figures S2D and 5D) was computed using R. All other statistical tests were performed in Python using SciPy and Statsmodels (*71*, *74*).

Distributions were first tested for normality (Shapiro-Wilk test) and homoscedasticity (Levene’s or Fligner Test) and then compared using the appropriate parametric or non-parametric tests, as applicable. In cases of multiple comparisons between the three developmental stages, distributions were first compared using ANOVA/ Kruskal Wallis (unpaired) or Friedman test (paired) with a significance threshold of 5% followed by appropriate post hoc pairwise comparisons with Holm-Bonferroni correction (*74*, *75*). Cohen’s d (effect size) was computed using Pingouin (*76*). Whiskers on the box plots represent 1.5 times the interquartile range (IQR).

## Acknowledgments

The authors would like to thank Prof. Masahiko Hibi and Prof. Koichi Kawakami for the transgenic lines used in this study and for the Tol2 transposase. Thanks to Dr. Aalok Varma, Mr. Sudeepta Sarkar, Dr. Anal Kumar, Dr. Anzal KS and Dr.

Nagaraju Dhanyasi for scientific and technical inputs. We would also like to thank Prof. Upi Bhalla and Dr. Baskar Bakthavachalu for insightful discussions. Further, we would like to thank the funding agencies listed below for support and Mr. P.T. Jagadeesh for the maintenance of our fish lines. We wish to thank Central Imaging and Flow facility (CIFF) - NCBS, CIFF - INSTEM, mechanical/electronics workshops and Sangers sequencing facilities for enabling this research.

## Funding

Wellcome Trust-DBT India Alliance Intermediate fellowship 500040/Z/09/Z (VT) Wellcome Trust-DBT India Alliance Senior fellowship IA/S/17/2/503297 (VT) Department of Biotechnology, Government of India BT/PR4983/MED/30/790/2012 (VT)

Department of Biotechnology, Government of India BT/PR51360/MED/122/373/2024 (VT)

Science and Engineering Research Board EMR/2015/000595 (VT) Department of Atomic Energy RTI 4006 (VT) and

NCBS-TIFR graduate student fellowships (SV, SP and SN).

## Author contributions

Conceptualization: SV, VT Methodology: SV, SN, SC, VT Investigation: SV, SC, SP Visualization: SV, SC, VT Funding acquisition: VT Project administration: VT Supervision: VT

Writing – original draft: SV, VT

Writing – review & editing: SV, SN, SC, VT

## Competing interests

Authors declare that they have no competing interests.

## Data and materials availability

All data are available in the main text or the supplementary materials. The raw data and scripts (used for experiment and analysis) will be made publicly available on Zenodo before final acceptance.

## Supplemental Materials

Figs. S1 to S5

Tables S1

Movies S1

## Notes

### Competing Interest Statement

The authors have declared no competing interest.

### Summary of Updates

Added a computational model Added data from two new experiments Rewrote parts of manuscript and changed title for better clarity

