## Supplementary Figures 1-5 and Supplementary Table 1 for "Developmental synaptic pruning sharpens neural representations for predictive motor control"

**Fig. S1.**

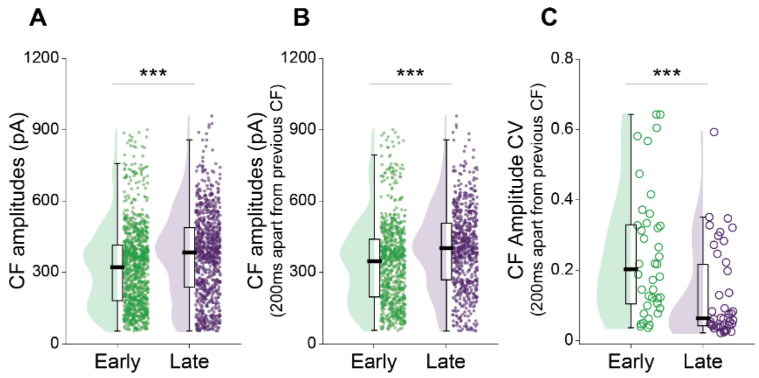


**Figure S1 Developmental changes in spontaneous CF inputs to PNs are independent of short-term depression**

**A.** Pooled amplitudes of all spontaneous CF EPSCs recorded from multiple PNs. Median ± IQR : 320.758 ± 234.027 (early), 383.16 ± 250.369 (late) (Mann-Whitney U test, P = 1.3e-12, Cohen's d = 0.294).

**B.** Pooled CF EPSC amplitudes that are spaced by at least 200 ms from the previous EPSC. Median ± IQR : 345.767 ± 243.107 (early), 401.269 ± 238.813 (late) (Mann-Whitney U test, P = 2.2e-14, Cohen's d = 0.356)

**C.** Coefficient of variation (CV) for spontaneous CF EPSCs that are spaced by at least 200 ms from the previous EPSC. Median ± IQR: 0.202 ± 0.225 (early), 0.063 ±0.174 (late) (Mann-Whitney U test, P = 0.0003, Cohen's d = 0.734).

Holding potential: -55mV. N = 40 and 42 cells (from 34 and 40 larvae) for early and late stages, respectively.
Colours *green* and *purple* represent early (4-5 dpf) and late (11-14 dpf) larval stages, respectively. ***P < 0.001.

---

**Fig. S2.**

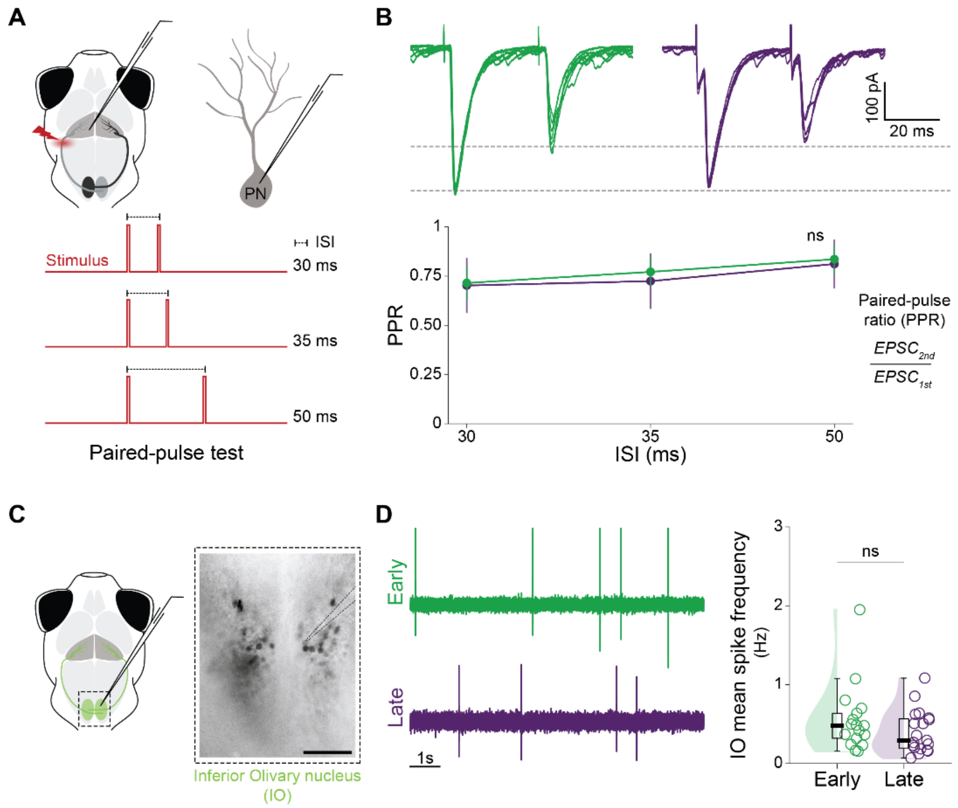


**Figure S2 Firing rate and release probability of presynaptic CFs do not change during the first two weeks of larval development.**

**A.** Top: Schematic of *in vivo* whole-cell patch-clamp recording from PNs with simultaneous paired-pulse stimulation of CFs using a bipolar electrode. Bottom: Interstimulus intervals (ISIs) of 30 ms, 35 ms, and 50 ms were used to assess the presynaptic release probability at the CF- PN synapses. Evoked CF EPSCs were recorded from PNs. Paired pulse ratio was calculated as the ratio of second to first evoked CF EPSC amplitude from 3-10 trials.

**B.** Top: Representative traces of CF EPSCs evoked by paired stimulus pulses at an ISI of 35 ms for early (left, green) and late (right, purple) larval stages (Holding potential: -55mV). Bottom: Paired Pulse Ratios (PPR) for early and late larval stages at 30 s, 35 s, 50 s. ISI significantly improved the model ($P < 0.001$) whereas no significant effect of developmental stage was seen on PPR ($P = 0.559$) using linear mixed-effects model with ANOVA. N = 10, 11 cells (from 10, 11 larvae) for early and late stages, respectively.

**C.** Left: Schematic of loose-patch recording from the soma of inferior olivary neurons in a ventral-up preparation. Right: Inferior olivary (IO) neurons tagged with green fluorescent protein in a transgenic hspGFFDMC28C:gal4; UAS:gfp larva. Targeted loose-patch recordings were performed from IO neurons. Scale bar 20 µm.

**D.** Left: Representative loose-patch recordings performed from early (top, green) and late (bottom, purple) stage larvae. Right: Mean spike frequency (Hz) for individual IO neurons does not change between early and late developmental stages. Median ± IQR : 0.48 ± 0.316 (early), 0.293 ± 0.371 (late) (Mann-Whitney U test, $P = 0.281$, Cohen's d = 0.433). N = 17, 19 cells (from 10, 8 larvae) for early and late stages, respectively.

Colours *green* and *purple* represent early (4-5 dpf) and late (11-14 dpf) larval stages, respectively. ns $P > 0.05$.

**Fig. S3.**

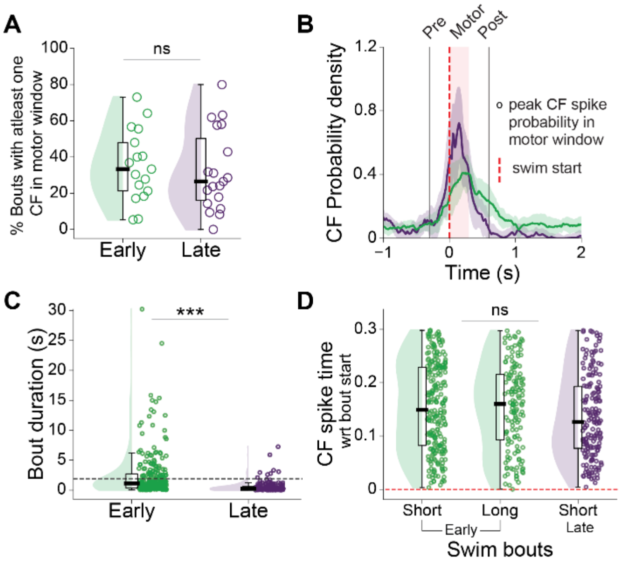


### Figure S3 Frequency of CF inputs to Purkinje neurons (PNs) at bout onset is independent of bout duration

**A.** Percentage of total swim bouts that resulted in at least one CF input during the 300 ms motor window (described in Fig. 3E (top)). Median ± IQR : 33.333 ± 26.723 (early), 26.47 ± 34.145 (late) (t-test, P = 0.672, Cohen's d = 0.142).

**B.** CF probability density around the start of swim bouts. Median ± IQR across cells is shown for early- and late- stage larvae, in contrast to Fig. 3D where data was pooled from all cells.

**C.** Durations of swim bouts that resulted in at least one CF input during the motor window. Median ± IQR : 1.117 ± 2.347 (early), 0.207 ± 0.47 (late) (Mann-Whitney U test, P = 5.5e-20, Cohen's d = 0.628). Black dashed line marks the threshold (calculated as the 95th percentile of bout durations in late stage larvae) for classification of bouts as short and long.

**D.** Timing of CF inputs recorded in PNs during the motor window for long and short bouts (Data pooled from swim bouts shown in panel C). Median ± IQR : 0.149 ± 0.146 (short bouts, early stage), 0.160 ± 0.122 (long bouts, early stage), 0.126 ± 0.116 (short bouts, late stage) (Kruskal Wallis test, P = 0.054).

N = 17, 19 cells (from 16, 19 larvae) for early and late stages, respectively. Colours *green* and *purple* represent early (4-5 dpf) and late (15-19 dpf) larval stages, respectively. Red dashed line indicates the onset of swim bouts. ***P < 0.001, ns P > 0.05.

---

**Fig. S4**

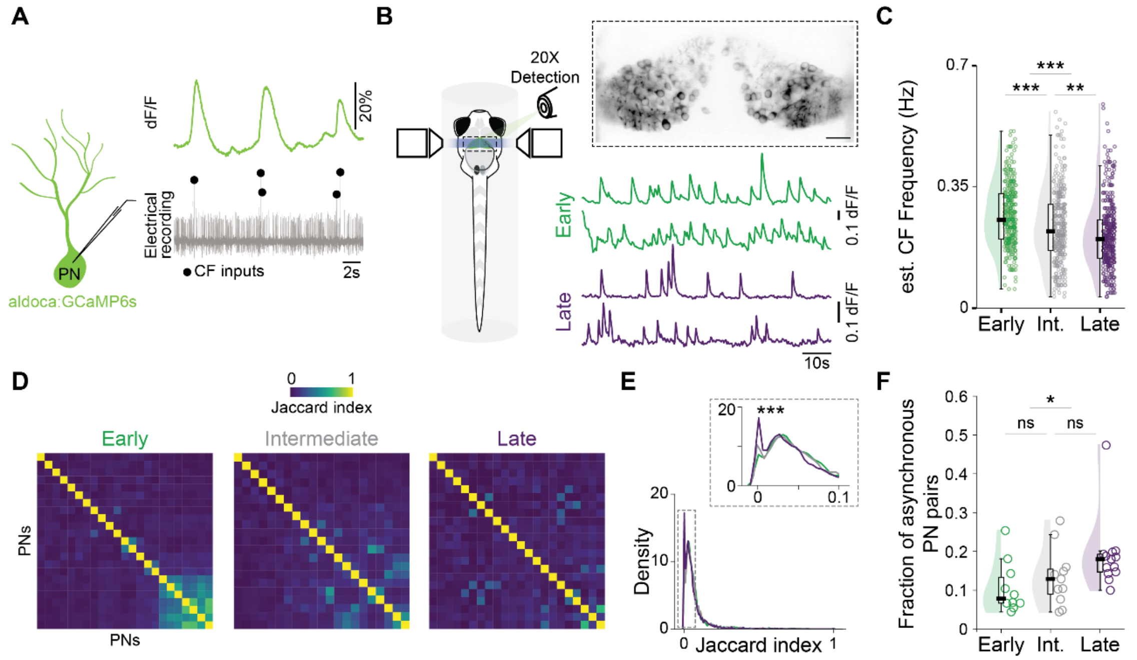


**Figure S4 Signatures of population wide specialization in Purkinje neurons (PNs) over early development.**

**A.** Representative widefield calcium imaging with simultaneous loose-patch recording from a PN showing large calcium transients associated with CF inputs. Transgenic aldoca:GCaMP6s; Nacre-/- larva was used to measure calcium activity in PNs.

**B.** Left: Schematic showing spontaneous light-sheet imaging of calcium activity in PNs. Right, top: Average GCaMP6s calcium activity of PNs in a single plane recording. Scale bar 20 µm. Right, bottom: Representative calcium activity traces from a pair of PNs in early and late stage larvae.

**C.** Decrease in estimated mean CF input frequency (Hz) over development. Median ± IQR : 0.255 ± 0.13 (early), 0.222 ± 0.133 (intermediate), 0.2 ± 0.111 (late) (One-way ANOVA for multiple comparisons (P = 3.5e-13) followed by pairwise t-test with Holm-Bonferroni correction for early vs intermediate (P = 3.8e-05, Cohen's d = 0.336), intermediate vs late (P = 0.002, Cohen's d = 0.232), early vs late (P = 1.7e-14, Cohen's d = 0.602)).

**D.** Jaccard indices for all pairs of PNs from a single lobe of a representative larva in each developmental group.

**E.** Distribution of Jaccard indices calculated from 90 s time series (pooled PN pairs from all larvae for each developmental group). Inset: X-axis zoomed-in from 0 to 0.1 to show an increased occurrence of completely asynchronous pairs in the late larval stage. Asynchronous PN pairs were defined as those with Jaccard index (JI) < 0.01 (One-way ANOVA, P = 7.5e-09).

**F.** Increase in fraction of asynchronous PN pairs over development. Median ± IQR : 0.085 ± 0.066 (early), 0.135 ± 0.064 (intermediate), 0.186 ± 0.045 (late) (Kruskal Wallis test for multiple comparisons (P = 0.018) followed by pairwise Mann-Whitney U test with Holm-Bonferroni correction for early vs intermediate (P = 0.378, Cohen's d = 0.385), intermediate vs late (P = 0.127, Cohen's d = 0.654), early vs late (P = 0.025, Cohen's d = 1.006)).

N = 326, 329, 381 cells (from 10, 11, 13 larvae) for early, intermediate and late stages, respectively. Colours *green*, *grey* and *purple* represent early (4-5 dpf), intermediate (7-8 dpf) and late (11-14 dpf) larval stages, respectively. ***P < 0.001, **P < 0.01, *P < 0.05, ns P > 0.05.

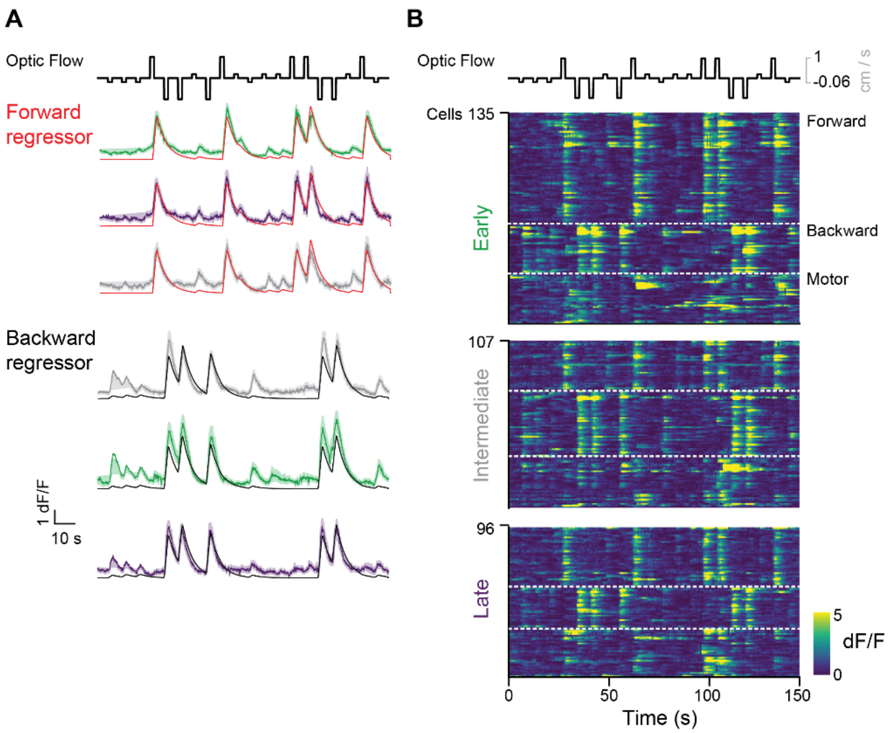


**Figure S5 Classification of Purkinje neurons (PNs) as forward-, backward- or motor-tuned using multiple linear regression.**

**A.** dF/F for PNs classified as forward (top) or backward (bottom) tuned (Mean ± SEM). A kernel with fast rise and slow decay was convolved with forward and backward optic flow to get the forward regressor (red, top) and backward regressor (black, bottom), respectively. For each larva, the swim speed trace was similarly convolved to get a motor regressor. With the help of these three regressors, multiple linear regression was performed on individual PN's dF/F trace. All cells with $R^2 \geq 0.4$ were classified as forward, backward or motor based on their largest regression coefficient.

**B.** Heat map of dF/F from all cells classified as forward, backward or motor (ordered from top to bottom, separated by white dashed lines) for early, intermediate and late larval stages.

Colours *green*, *grey* and *purple* represent early (4-5 dpf), intermediate (7-8 dpf) and late (11-14 dpf) larval stages, respectively.

---

### Table S1. Parameters for extraction of raw GCaMP6s fluorescence from PNs

Parameters used for extraction of GCaMP6s fluorescence signal from cells identified in suite2P. Left : Light-sheet images (1P). Right : two-photon images.

| ops_params | values_1P | values_2P |
|---|---|---|
| suite2p_vers | 0.14.2 | 0.14.2 |
| look_one_lev | 0 | 0 |
| fast_disk | [] | [] |
| delete_bin | FALSE | FALSE |
| mesoscan | FALSE | FALSE |
| bruker | FALSE | FALSE |
| bruker_bidire | FALSE | FALSE |
| h5py | [] | [] |
| h5py_key | data | data |
| nwb_file | | |
| nwb_driver | | |
| nwb_series | | |
| save_path0 | | |
| save_folder | [] | [] |
| subfolders | [] | [] |
| move_bin | FALSE | FALSE |
| nplanes | 1 | 1 |
| nchannels | 1 | 1 |
| functional_cl | 1 | 1 |
| tau | 1.5 | 1.5 |
| fs | 25 | 8 |
| force_sktiff | FALSE | FALSE |
| frames_inclu | -1 | -1 |
| multiplane_p | 0 | 0 |
| ignore_flyba | [] | [] |
| preclassify | 0 | 0 |
| save_mat | FALSE | FALSE |
| save_NWB | 1 | 1 |
| combined | 1 | 1 |
| aspect | 1 | 1 |

| ops_params | values_1P | values_2P |
|---|---|---|
| do_bidiphase | FALSE | FALSE |
| bidiphase | 0 | 0 |
| bidi_correcte | FALSE | FALSE |
| do_registrati | 1 | 1 |
| two_step_re | 0 | 0 |
| keep_movie_ | TRUE | TRUE |
| nimg_init | 300 | 300 |
| batch_size | 250 | 1000 |
| maxregshift | 0.1 | 0.1 |
| align_by_cha | 1 | 1 |
| reg_tif | TRUE | TRUE |
| reg_tif_chan | FALSE | FALSE |
| subpixel | 10 | 10 |
| smooth_sign | 0 | 0 |
| smooth_sign | 4 | 4 |
| th_badframe | 1 | 1 |
| norm_frame | TRUE | TRUE |
| force_refImg | FALSE | FALSE |
| pad_fft | FALSE | FALSE |
| nonrigid | TRUE | TRUE |
| block_size | [128, 128] | [64, 64] |
| snr_thresh | 1.5 | 1.2 |
| maxregshiftN | 8 | 5 |
| 1Preg | TRUE | FALSE |
| spatial_hp_r | 42 | 42 |
| pre_smooth | 0 | 0 |
| spatial_tape | 40 | 40 |
| roidetect | TRUE | TRUE |
| spikedetect | FALSE | FALSE |
| sparse_mode | TRUE | TRUE |

| ops_params | values_1P | values_2P |
|---|---|---|
| spatial_scale | 0 | 0 |
| connected | TRUE | TRUE |
| nbinned | 5000 | 5000 |
| max_iteratio | 20 | 20 |
| threshold_sc | 1 | 1 |
| max_overlap | 0 | 0 |
| high_pass | 6 | 100 |
| spatial_hp_d | 25 | 25 |
| denoise | 0 | 0 |
| anatomical_ | 3 | 3 |
| diameter | 20 | 12 |
| cellprob_thre | 0 | 0 |
| flow_thresho | 2 | 2 |
| spatial_hp_c | 0 | 0 |
| pretrained_n | cyto | cyto |
| soma_crop | 1 | 1 |
| neuropil_ext | TRUE | TRUE |
| inner_neurop | 10 | 6 |
| min_neuropi | 350 | 200 |
| lam_percent | 50 | 50 |
| allow_overla | FALSE | FALSE |
| use_builtin_ | FALSE | FALSE |
| classifier_path | | |
| chan2_thres | 0.65 | 0.65 |
| baseline | maximin | maximin |
| win_baseline | 60 | 60 |
| sig_baseline | 10 | 10 |
| prctile_basel | 8 | 8 |
| neucoeff | 0.7 | 0.7 |
| input_forma | tif | tif |

### Movie S1. Desynchronization of PNs during early larval development

Late-stage PNs exhibit more asynchronous calcium activity (right), than the early-stage (left). A subset of PNs from a single lobe of the representative larvae mentioned in Fig. S4D are shown. Transgenic aldoca:GCaMP6s; nacre -/- larvae were imaged at 25 Hz using light-sheet microscopy. Scale bar 20 µm.
